# Tumor-immune trajectory context connects static tissue architecture to clinical outcomes

**DOI:** 10.64898/2026.03.26.714521

**Authors:** Eric M. Cramer, Randy Heiland, Heber Lima da Rocha, Daniel R. Bergman, Joe W. Gray, Gordon B. Mills, Elana J. Fertig, Paul Macklin, Laura M. Heiser, Young Hwan Chang

## Abstract

Multiplexed tissue imaging (MTI) has revealed recurrent tumor microenvironment (TME) architectures with prognostic value, yet these measurements are inherently static, obscuring dynamic changes in the TME that govern therapeutic response. Here, we introduce a trajectory-centric framework that reconstructs continuous TME dynamics by integrating agent-based mathematical modeling and simulation with state space analysis. This approach yields a mechanistically constrained reference landscape built entirely from *in silico* simulation, and onto which static patient biospecimens can be projected and mapped onto simulated TME trajectories. Systematic simulation of tumor-immune interactions in triple-negative breast cancer identifies six metastable TME states connected by transition pathways spanning immune control to immune escape. Mapping MTI data from two independent patient cohorts, including longitudinal samples from a randomized immunotherapy trial, validates this landscape by positioning individual biospecimens along inferred TME trajectories rather than in static states. We show that treatment-phase TME states, but not pre-treatment configurations, robustly predict immunotherapy response, and identical terminal states can arise from distinct trajectory histories corresponding to immune failure or resolved inflammation. Thus, this framework enables mechanistic simulations to define a reference dynamical landscape that serves as a coordinate system for interpreting static clinical spatial data, providing a principled basis for evaluating consistency, predictiveness, and clinical relevance across independent patient cohorts. Altogether, this study advances spatial tumor profiling from static state classification of human tissues to dynamic trajectory inference, establishing a quantitative framework for trajectory-informed, state-guided, and temporally adaptive immunotherapy strategies.

## Introduction

Tumors are evolving multicellular ecosystems with compositions, spatial organizations, and functional interactions that continuously change over time.^1,2^ In the tumor microenvironment (TME), cancer cells, immune populations, stromal elements, and vasculature dynamically interact through feedback loops that determine immune efficacy, immune escape, disease progression, and therapeutic response.^3,4^ Multiplexed tissue imaging (MTI) has substantially expanded our ability to characterize the TME at single-cell resolution while preserving spatial context,^5–7^ revealing that tumors self-organize into recurring spatial and cellular configurations (often referred to as archetypes, ecotypes, or niches) which associate with clinical outcomes across cancer types.^8–11^ In triple-negative breast cancer (TNBC), for example, architectures of immune infiltration and measures of cell-cell interactions between tumor and immune cells have been linked to prognosis and immunotherapy response.^8,11,12^ Yet a fundamental challenge arises from the invasive nature of clinical tissue sampling: MTI data necessarily capture static snapshots of a dynamic system.^5^ Even longitudinally collected serial samples represent discrete observations separated by unobserved biological evolution rather than continuous temporal trajectories. Consequently, current spatial analysis approaches primarily interpret biological meaning from static configurations, leaving a critical question: how do organizational structures within the TME arise, persist, and transition over time?^13,14^

This inability to fully measure the spatiotemporal changes in cellular and molecular landscapes of the TME is particularly consequential for interpreting responses to anti-tumor therapies such as immunotherapy. Clinical response reflects both the baseline TME composition and the change in tumor-immune interactions following therapeutic perturbation.^15,16^ For example, both a tumor with failed immune engagement and one reflecting a resolved inflammatory contexture may look identical in a static measurement of an instantaneous moment even though they have vastly different histories and futures in their immune state. Without trajectory context, such distinctions are invisible, confounding biomarker development and therapeutic decision-making.

Several new techniques have begun to address this temporal gap. Mathematical frameworks have been proposed to infer cell type population-level dynamics from spatial snapshots, demonstrating that local neighborhood structure and proliferation markers can encode information about underlying tissue dynamics.^14,17^ One such approach, agent-based modeling, has emerged as a powerful mathematical tool for simulating multicellular systems. Agent-based modeling takes a bottom-up approach to simulating the TME by representing each cell as an autonomous "agent" that follows biologically motivated rules governing cell behavior, communication, and fate decisions.^18–21^ Notably, recent developments to the PhysiCell agent-based modeling framework enable human-interpretable *rules* that are standardized logical formulations mapping environmental signals to cellular behaviors, thus enabling rigorous mechanistic definition of cell-cell interactions.^22^ These rules follow the flow of if-then statements, such as **if** oxygen concentration is sufficient **then** tumor cells are less likely to undergo necrosis. We hypothesize that if the mechanistic structure imparted by an ABM is fundamentally sound, systematic perturbation of the model’s parameters can successfully capture and encode the diverse biological heterogeneity observed in clinical data. However, a unifying framework that connects mechanistic simulations such as agent-based modeling to clinical spatial data and enables patient samples to be interpreted within the context of tumor microenvironment evolution has remained elusive.^22–26^

Here, we introduce a computational framework that uses a mechanistic agent-based model (ABM) to reconstruct continuous TME dynamics, generating a mechanistically constrained reference landscape onto which static MTI samples can be mapped – empowering inference of TME evolution from individual patient samples. We demonstrate this approach using MTI-profiled biospecimens from two independent cohorts of patients with TNBC. Crucially, this landscape is computed once and serves as a permanent reference coordinate system: any new patient sample can be positioned within it without rerunning the ABM or recomputing the landscape. Analysis of this landscape identifies six canonical TME states spanning favorable Effector-Dominant to unfavorable Immune-Excluded tissue architectures. Mapping longitudinal samples from a randomized clinical trial onto the landscape reveals that treatment-phase TME states rather than initial pre-treatment configurations robustly predict immunotherapy response. Finally, by summarizing the ABM dynamics with a Markov state model of transitions among these six TME states, we simulate therapeutic interventions and identify CD8^+^ T cell exhaustion-driven transition points that define windows for potentially effective immune modulation. Our framework reframes spatial tumor profiling from a static classification problem to a trajectory-centric paradigm, providing a quantitative foundation for TME state-guided therapeutic strategies.

## Results

### Agent-based modeling reconstructs continuous TME dynamics from static tissue snapshots

In clinical practice, biospecimens are collected at non-uniform intervals across patients, often weeks, months or even years apart, obscuring the dynamic processes that govern treatment response.^5,27,28^ To address this fundamental limitation, we adapted a previously published, mechanistic ABM of tumor-immune cell interactions that captures key cellular populations and biological processes governing immune activation and suppression (**Fig. 1A**).^22^ Specifically, the model simulates how cytokine-driven macrophage polarization and T cell exhaustion create a transition from an initial anti-tumor response to an immunosuppressive environment that permits tumor regrowth.

**Figure 1.**
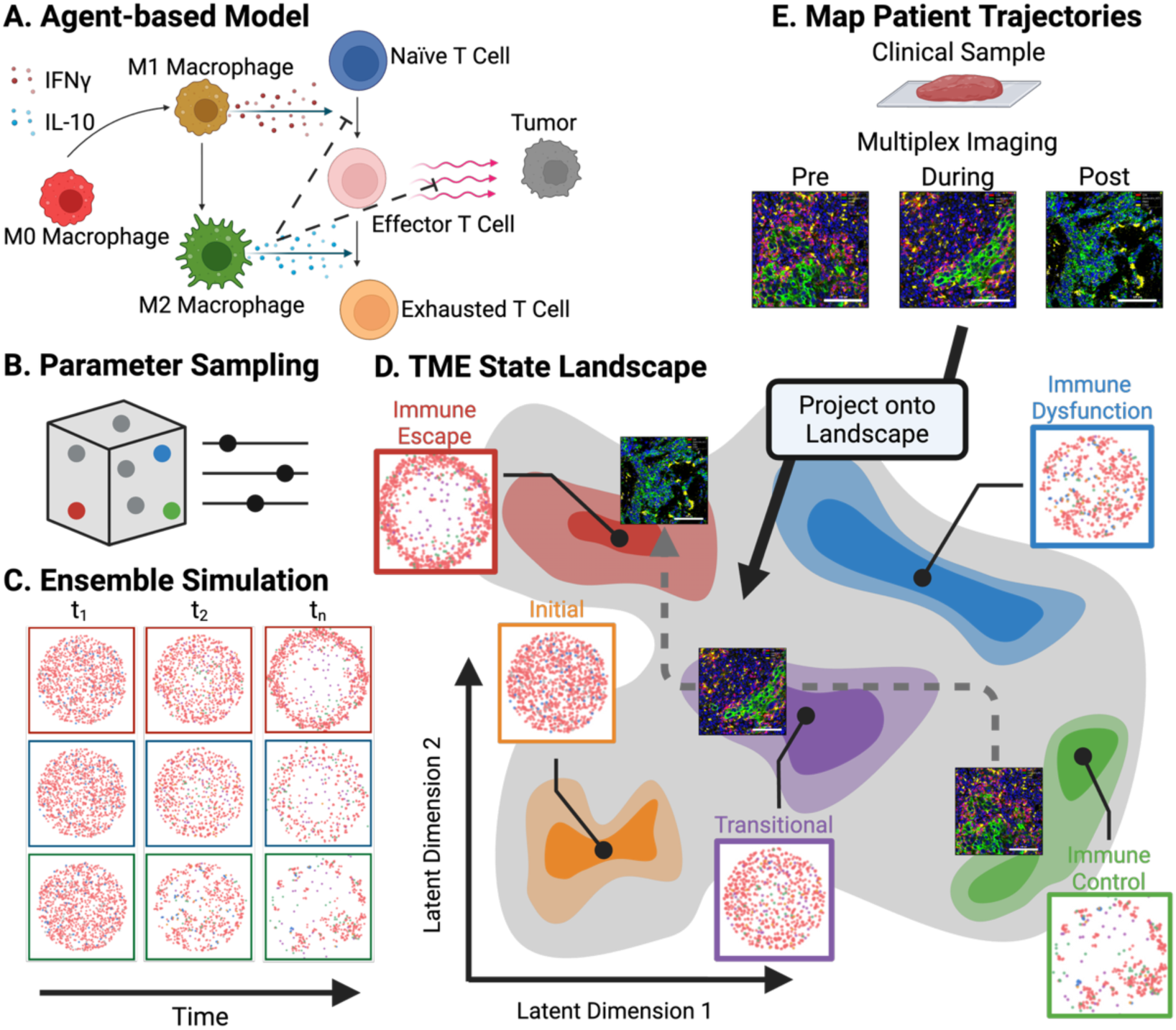
Agent-based modeling enables temporal reconstruction of tumor microenvironment dynamics from static imaging data. **A,** Agent-based model (ABM) of tumor-immune cell interactions in the TME adapted from Johnson et al. *Cell* 2025 **B,** Parameters are sampled from a Latin Hypercube to generate different input parameter value combinations. **C,** Simulation of the ABM with the many different input parameter value combinations to form an “ensemble” of simulated TME trajectories. **D,** Clustering the ensemble of trajectories forms a TME state landscape with regions of where simulations possessed similar or dissimilar spatial configurations of cells for discrete periods of time. Simulated tumors begin in an initial state and proceed to subsequent states based on the effects of their input parameter values analogous to a Waddington landscape^33^ for tissue. **E,** Multiplexed imaging of clinical samples can be projected or “mapped” onto the pre-computed TME state landscape to contextualize potential tumor trajectories.

We systematically explored the parameter space of the ABM to capture the diversity of tumor-immune behaviors that can arise from shared biological rules (**Fig. 1B**, see Methods: Parameter Sampling). This approach allows the ABM to define a global dynamical landscape that reflects inter- and intra-patient heterogeneity in tumor-immune dynamics rather than a single representative trajectory. To this end, we generated ABM parameter sets using Latin hypercube sampling (**Fig. 1B, Supplementary Table 1**) and ran simulations with each parameter set from three randomized initial cell positions (**Fig. 1C**). Initial cell positions represented mixing of tumor and immune cells consistent with tumor growth. Cell positions were constrained to preserve comparable spatial organization, ensuring that trajectory differences reflected parameter effects rather than spatial artifacts introduced by initial conditions (**Supplementary Fig, 1a-c**). Simulations were run forwards over two weeks of biological time assuming a therapeutic or evolutionary perturbation (represented by the parameter sample from the Latin Hypercube). We generated dense, multivariate time-series data describing the state of each simulation at each time point by calculating spatial statistics characterizing cell type composition, cell connectivity graph network measures, and cell-cell interactions – features directly analogous to those extracted from MTI data (see Methods: Spatial Statistics Calculation).

We then transformed these multivariate time series using time-delay embedding based on Takens’ theorem ^29–32^, generating a representation of the data that encodes system dynamics by concatenating sliding temporal windows of timepoints into single feature vectors for each simulation (**Supplementary Fig. 2**). This embedding has been shown to preserve the topology an underlying dynamical system, and in our study defines a landscape of TME tissue-level dynamics analogous to a Waddington landscape for individual cells.^29,30,33^ Clustering the time-delay embedding identified discrete regions of the landscape where simulations possessed similar spatiotemporal configurations of cells for discrete periods of time – or metastable “TME states” that the simulated tumor tissue occupy along their various trajectories (**Fig. 1D**). The TME state landscape generation procedure is conceptually illustrated in more detail in **Supplementary Fig. 3a-b**. Computing this landscape *a priori* from spatial measurements enables future projection or mapping of clinical samples onto the TME state landscape (**Fig. 1E**).

### ABM-derived state landscape encodes biologically interpretable temporal dynamics

Our clustering of the time-delay embedding resolved six discrete TME states in this landscape that we sought to characterize (**Table 1, Supplementary Fig. 3C**). Analysis of all measured simulation features indicated states of the TME state landscape are distinguished by coherent and biologically interpretable signatures (**Fig. 2A**). The **Effector-Dominant** (State 1; Eff⁺/Exh⁻; Green) represents active immune engagement, characterized by abundant functional effector CD8⁺ T cells and relatively lower exhausted CD8^+^ T cells. This state exhibits increased M1 macrophage connectedness and interaction with each other and tumor cells, along with elevated M1-M2 macrophage ratios indicative of pro-inflammatory microenvironments. Notably, terminal Effector-Dominant equilibrium simulations show decreased tumor cell proportions alongside sustained effector CD8^+^ T cell presence, consistent with ongoing immune-mediated tumor control. **Undifferentiated** (State 2; Eff⁰/Exh⁰; Orange) occupies the origin of the TME state landscape and represents an uncommitted configuration in which immune populations have not yet polarized toward effector vs. exhausted or pro-inflammatory vs. anti-inflammatory differentiation programs and tumor content is high. **Pre-Exclusion Transitional** (State 3; Eff↓/Exh↑; Sage) represents a branch point for trajectories that fail to achieve or sustain effector CD8^+^ T cell activation. **Exhausted-Dominant** (State 4; Eff⁻/Exh⁺; Grey) is a terminal state with CD8^+^ T cells present but dysfunctional, having lost effector capacity. This state exhibits elevated centrality of exhausted CD8⁺ T cells and enrichment of M2-polarized macrophages, consistent with immune dysfunction and a pro-tumorigenic TME despite immune infiltration. **Pre-Exhaustion Transitional** (State 5; Eff⁻/Exh↑; Purple) represents committed decline toward CD8^+^ T cell exhaustion, while **Immune-Excluded** (State 6; Eff⁻/Exh⁻; Gold) corresponds to classical "cold" TMEs with tumor dominance characterized by high tumor centrality and minimal CD8⁺ T cell presence. By explicitly separating effector status (Eff⁺/⁻) from exhausted status (Exh⁺/⁻), this nomenclature extends conventional hot/cold classifications and distinguishes infiltrated but dysfunctional TMEs that are biologically and therapeutically distinct.

**Figure 2.**
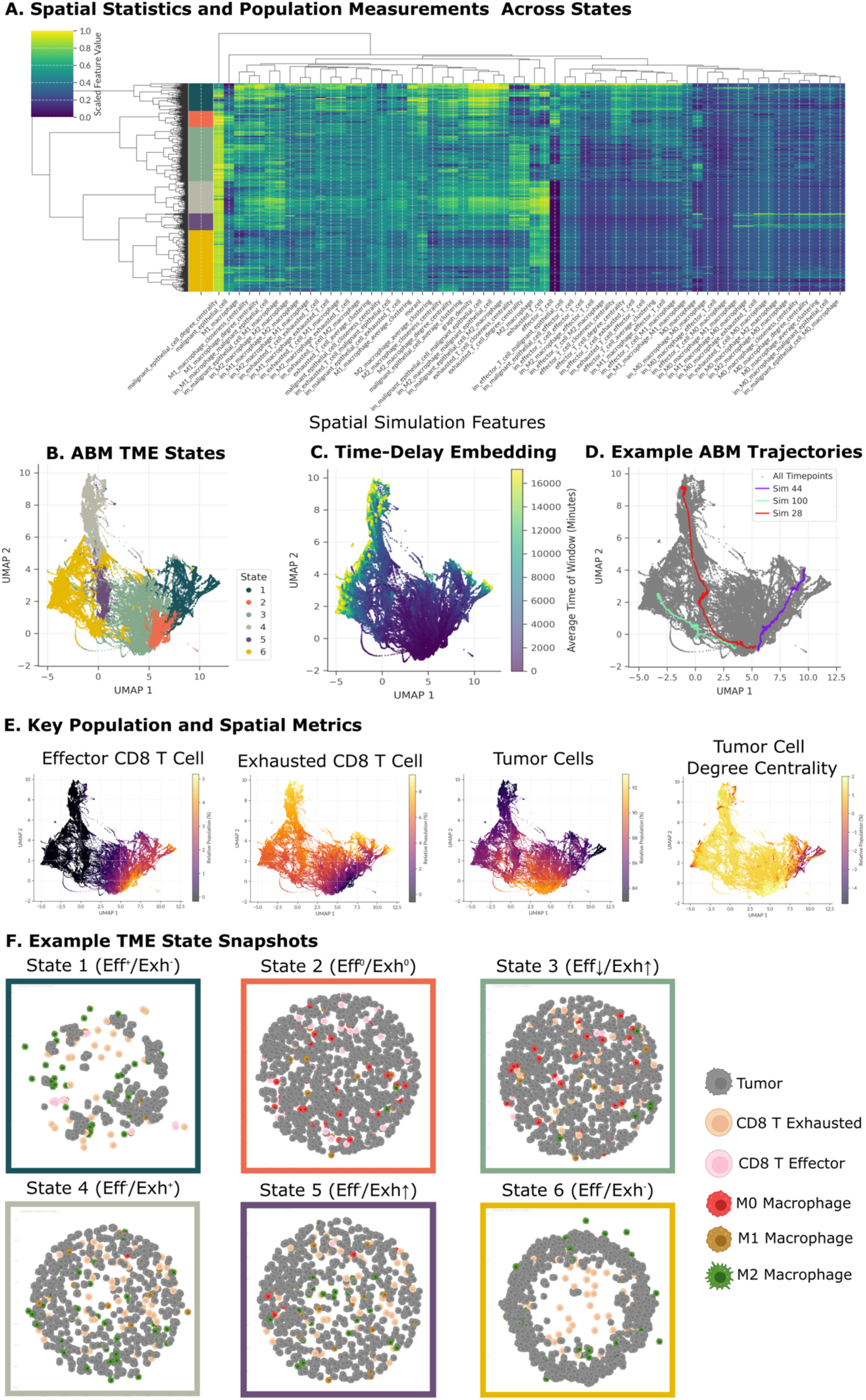
The ABM-derived TME space captures temporal dynamics and defines biologically interpretable TME states. **A,** Clustermap of spatial and population features across Leiden communities grouped by state assignment. Distinct profiles distinguish origin, transitional, and terminal configurations. **B,** Representative simulation snapshots for each TME state, illustrating characteristic architectures: active effector immune infiltration and activity (State 1), mixed infiltration (State 2), early immune decline (State 3), exhausted but infiltrated (State 4), committed decline to exhaustion phenotype (State 5), and immune excluded (State 6). **C,** Biological characterization of state landscape. UMAP projections show continuous variation in effector CD8⁺ T cell proportion, exhausted CD8⁺ T cell proportion, tumor cell degree centrality, and tumor cell proportion across states. **D,** UMAP visualization of time-delay embedded simulation data, colored by average time of timesteps in window. **E,** TME state assignments from hierarchical clustering. Six states defined: Effector-Dominant (State 1, Eff^+^/Exh^-^), Undifferentiated (State 2, Eff^0^/Exh^0^), Early-Transitional (State 3, Eff^,i.^/Exh^τ^), Exhausted-Dominant (State 4, Eff^-^/Exh^+^), Late-Transitional (State 5, Eff^-^ /Exh^τ^), Immune-Excluded (State 6, Eff^-^/Exh^-^). **F,** Trajectory structure of the state landscape. Colored lines trace three representative simulations from origin area (Undifferentiated, State 2) through divergent paths.

**Table 1.** TME State Nomenclature and Characteristics.

| State |  | Nomenclature | Descriptive Features | Biological Interpretation |
| --- | --- | --- | --- | --- |
| 1 |  | Effector-Dominant (Eff<sup>+</sup>/Exh<sup>-</sup>) | High effector CD8 T cell density and activity; organized immune-tumor interface with niches for effector CD8 T cell maintenance | Active anti-tumor immunity with structured immune response; T cells organized at tumor margin |
| 2 |  | Undifferentiated (Eff<sup>0</sup>/Exh<sup>0</sup>) | Balance of effector and exhausted CD8 T cells with tumor-immune mixing; macrophage polarization skews towards M1 | Active immune-tumor contact with mounting exhausted immune response |
| 3 |  | Pre-Exclusion Transitional (Eff<sup>↓</sup>/Exh<sup>↑</sup>) | Reduced effector immune presence; macrophage polarization ratio skews towards M1 (pro-inflammatory) | Metastable state between immune engagement and exclusion; potentially susceptible to perturbation |
| 4 |  | Exhausted-Dominant (Eff<sup>-</sup>/Exh<sup>+</sup>) | High exhausted CD8 T cell density; macrophage polarization almost exclusively M2 (anti-inflammatory) | Immunosuppressive microenvironment has developed preventing tumor control |
| 5 |  | Pre-Exhaustion Transitional (Eff<sup>-</sup>/Exh<sup>↑</sup>) | Balance of effector CD8 T cell to exhausted CD8 T cell skews towards exhausted; macrophage-dominant infiltrate skews towards M2 polarization; tumor population transiently checked yet uncontrolled | Immunosuppressive microenvironment emerging; potential myeloid-mediated effector CD8 T cell exclusion |
| 6 |  | Immune-Excluded (Eff<sup>-</sup>/Exh<sup>-</sup>) | Minimal immune infiltration; low T cell connectivity; absence of immune-preserving niches; Macrophage population exclusively M2 polarized; tumor cell compaction | Failed immune recognition or complete immune evasion |

Six canonical TME states derived from agent-based modeling of tumor-immune dynamics. Color swatches indicate the visualization scheme used throughout this manuscript.

We sought to visualize the high-dimensional TME state landscape encoded by the time-delay embedding to evaluate the distribution and relationships between key features in the TME state landscape. Visualization of the time-delay embedding in two dimensions using UMAP, where each point represents an embedded time window, revealed the distribution of key features. Coloring the embedding according to the cluster label (**Fig. 2B**) and the maximum simulation time (**Fig. 2C**) of each embedded window revealed a clear temporal organization: early time windows cluster near a common origin state while later windows progressively diverge, reaching terminal states toward the periphery. Plotting the time-delay embeddings of individual simulation trajectories on top of the overall time-delay embedding confirmed that simulations originate from a shared region of the TME state landscape before branching outwards along distinct paths as the simulations progress (**Fig. 2D**). This branching pattern indicates that forcing similarity between the spatial statistics of the initial condition replicates used by different simulation runs of the ABM successfully controlled for starting states. Furthermore, the branching pattern demonstrated that the time-delay embedding captures temporal progression of simulations rather than static similarity alone.

We selected key cellular population and spatial statistic features (**Fig. 2A**) that varied dramatically between TME states to visualize further on the TME state landscape (**Fig. 2E**). Effector and exhausted CD8⁺ T cell proportions define opposing gradients across the TME state landscape, capturing the transition from engagement of immune effectors to immune exhaustion. Tumor cell degree centrality and tumor cell proportion together reflect increasing tumor dominance across unfavorable states. Representative simulation snapshots (**Fig. 2F**) provided qualitative confirmation of the differences between the distinct multicellular architectures underlying each state, ranging from the Effector-Dominant configurations with dispersed tumor regions to Immune-Excluded configurations dominated by dense, contiguous tumor masses.

### Clinical MTI data map coherently onto the ABM-derived TME state landscape

To test whether the computationally derived state landscape captures biologically and clinically relevant TME variation, we mapped patient samples from two independent TNBC cohorts into the coordinate system of the TME state embedding defined by the ABM (**Fig. 3A**). We selected datasets to explicitly compare the impacts of mapping MTI datasets to our map in cases with single snapshot and longitudinal data tracing patient response. The first dataset comprises Multiplexed Ion Beam Imaging (MIBI) imaging of archival FFPE tissue microarray cores from 41 patients collected at a single snapshot in time from surgical resection biospecimens obtained between 2002 and 2015.^8^ The second dataset comprises longitudinal imaging mass cytometry (IMC) from the NeoTRIP randomized clinical trial (NeoTRIPaPDL1; NCT02620280), a multicenter, open-label Phase III study of neoadjuvant therapy for early high-risk and locally advanced TNBC.^11,27^ In this study, a total of 279 patients were randomized to chemotherapy alone (C; carboplatin and nab-paclitaxel, n = 141) or chemotherapy plus concurrent anti-PD-L1 immunotherapy (C & I; carboplatin, nab-paclitaxel, and atezolizumab, n = 138), with both agents administered together from day 1 of treatment—not sequentially. All drugs were given on day 1 of each 3-week cycle (carboplatin and nab-paclitaxel also on day 8) for eight cycles (∼24 weeks total). Tumor biospecimens were collected at three timepoints from 279 patients: baseline (pre-therapy), early on-treatment (day 1 of cycle 2, ∼3 weeks), and post-treatment (surgical excision immediately following the final cycle, ∼24 weeks). Pathological complete response (pCR; absence of invasive cells in breast and lymph nodes at surgical excision) served as the primary surrogate endpoint for correlative analyses.

**Figure 3.**
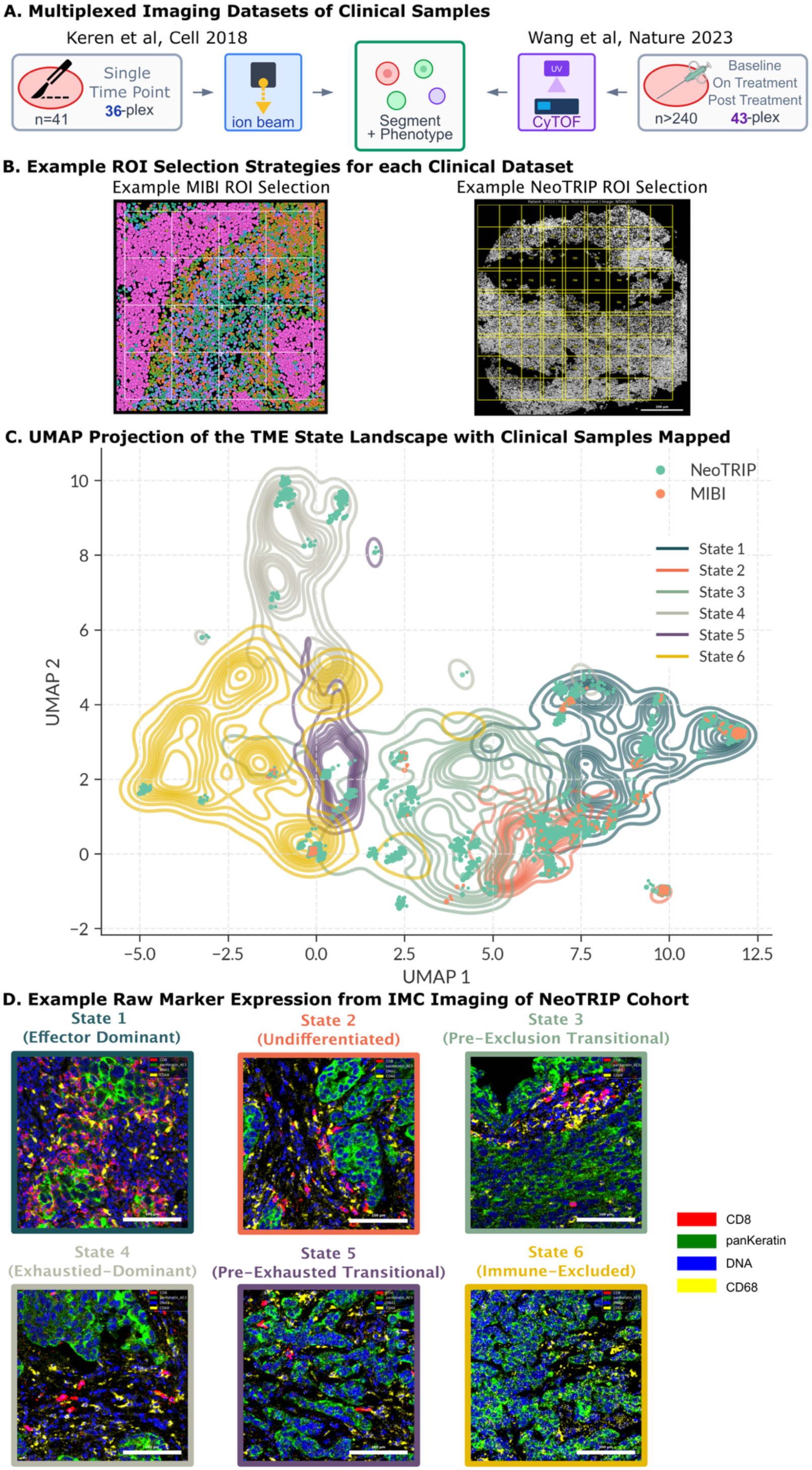
Mapping multiplexed imaging data from two independent TNBC cohorts onto the ABM-derived TME state landscape. **A,** Clinical datasets for validation. Left: MIBI-TOF data from Keren et al. (41 patients, single timepoint at surgical resection). Right: IMC data from NeoTRIP trial produced by Wang et al. (243 patients, longitudinal sampling at baseline, on-treatment, and post-treatment). Both modalities provide segmented single-cell data with spatial coordinates for state landscape mapping. **B,** Exemplars of ROI sampling strategy. Left: example taken from NeoTRIP data. Right: example taken from MIBI data. **C,** Projection of clinical ROIs onto ABM-derived TME state landscape. Contours show density of simulated time windows; points show MIBI cohort (orange) and NeoTRIP cohort (green) ROIs. Points dodged ≤0.2 units to facilitate visualization. **D,** Representative IMC images from the NeoTRIP cohort showing spatial signal intensity from ROIs assigned to each TME state. Markers shown: CD8, panKeratin, DNA/DAPI, and CD68.

We next investigated how MTI profiling of real human tumor tissues occupy regions of the TME state landscape. Both MTI data sets included spatially resolved cell phenotype labels and cell-level marker expression data which we used to refine the cell phenotype labels to harmonize with those of our ABM (**Supplementary Fig. 4**). After harmonization, we calculated the same population measures and spatial features as those we calculated at each ABM time step to facilitate mapping the biological samples onto the TME state landscape.

Spatial statistics (particularly graph based network measures such as centrality scores), however, are sensitive to the size and shape of the region being measured,^34^ and direct comparison across heterogeneous fields of view can obscure biological signal. To enable meaningful comparison between biological samples and ABM simulations, we addressed the scale dependence of graph-based spatial metrics often used in TME analysis by implementing a region-of-interest (ROI) sampling strategy.^34–36^ Our ROI sampling strategy (**Fig. 3B**, see Methods) tiles each clinical specimen with ROIs to best match the spatial scale and cell-connectivity graph density of ABM simulation snapshots. This approach recovers the full spectrum of local TME configurations within each sample while minimizing scale-induced bias (**Supplementary Fig. 5A-G**).

Neither MTI dataset measured any protein markers that would capture macrophage polarization. As re-running the ABM simulations without macrophages would fundamentally alter the spatial relationships and cell-cell interactions formed within the ABM output, we instead chose to assess the ability to map human tumor samples into the TME state landscape without macrophage polarization information; the presence of macrophages in the patient samples would be implicitly captured by the spatial statistics measuring how the different T cell populations in our model form interactions with the tumor cells. We tested the procedure by mapping randomly selected ABM time windows back to the TME state landscape after removing their macrophage-related features (see Methods: State Landscape Mapping). We found state assignment for the tested time windows was preserved with 97% accuracy (**Supplementary Fig. 6A-B**). This internal validation supports the validity of the ABM-derived TME state landscape as a reference for clinical data lacking complete feature coverage.

Projection of the sampled clinical ROIs onto the validated TME state landscape revealed marked differences between cohorts (**Fig. 3C**). Mapped states correspond to recognizable biological phenotypes rather than arbitrary computational groupings (**Fig. 3D**). ROIs from the MIBI cohort were concentrated primarily in Effector-Dominant (State 1) and Undifferentiated (State 2), with no representation in Exhausted-Dominant (State 4). In contrast, ROIs from the NeoTRIP cohort spanned all six states. This difference was driven in part by differences in the prevalence of exhausted CD8+ T cells across datasets (**Supplementary Fig. 7A-C**). ROIs assigned to Effector-Dominant (State 1) exhibited dense CD8⁺ T cell infiltration alongside tumor cells, consistent with active immune engagement. In contrast, ROIs assigned to Immune-Excluded (State 6) showed sparse CD8⁺ T cell presence and tumor dominance, matching the "effector-cold" phenotype predicted by the simulated landscape. Consistent with the difference in the mapping between the two cohorts, the MIBI cohort had low abundance of exhausted

CD8+ T cells, while the exhausted CD8+ T cell population grows over the course of treatment in the NeoTRIP trial (**Supplementary Fig. 7A**). Together, these results demonstrate that clinical TNBC samples occupy structured, biologically interpretable regions of the ABM-derived TME state landscape, supporting its utility as a mechanistically grounded reference framework for integrating and biologically contextualizing clinical MTI data.

### ABM-derived TME states recapitulate spatial mixing phenotypes while improving prognostic resolution

Having established that clinical samples map coherently onto the TME state landscape, we next examined whether the ABM-derived TME state assignments simply recapitulate previously defined spatial phenotypes or instead provide additional clinically relevant information. We focused on the Keren/MIBI cohort, in which prior work defined a tumor-immune mixing score based on expert-curated spatial organization and demonstrated its association with disease-free survival (DFS).^8,37^

Each patient’s sample from the MIBI cohort was sub-sampled into ROIs (see Methods – ROI Sampling Strategy), which were independently mapped to the ABM derived TME state landscape (**Fig. 4A**). ROI-level assignments revealed substantial within-patient heterogeneity, consistent with spatially complex tumor-immune organizations. We obtained a patient-level measure estimating the dominant TME state per patient by aggregating ROI-level assignments and assigning the patient a state based upon their most frequent ROI state assignment. This patient-level summary of the states in our model enabled direct comparison of our simulated states with the patient-level mixing score classification from Keren et al.

**Figure 4.**
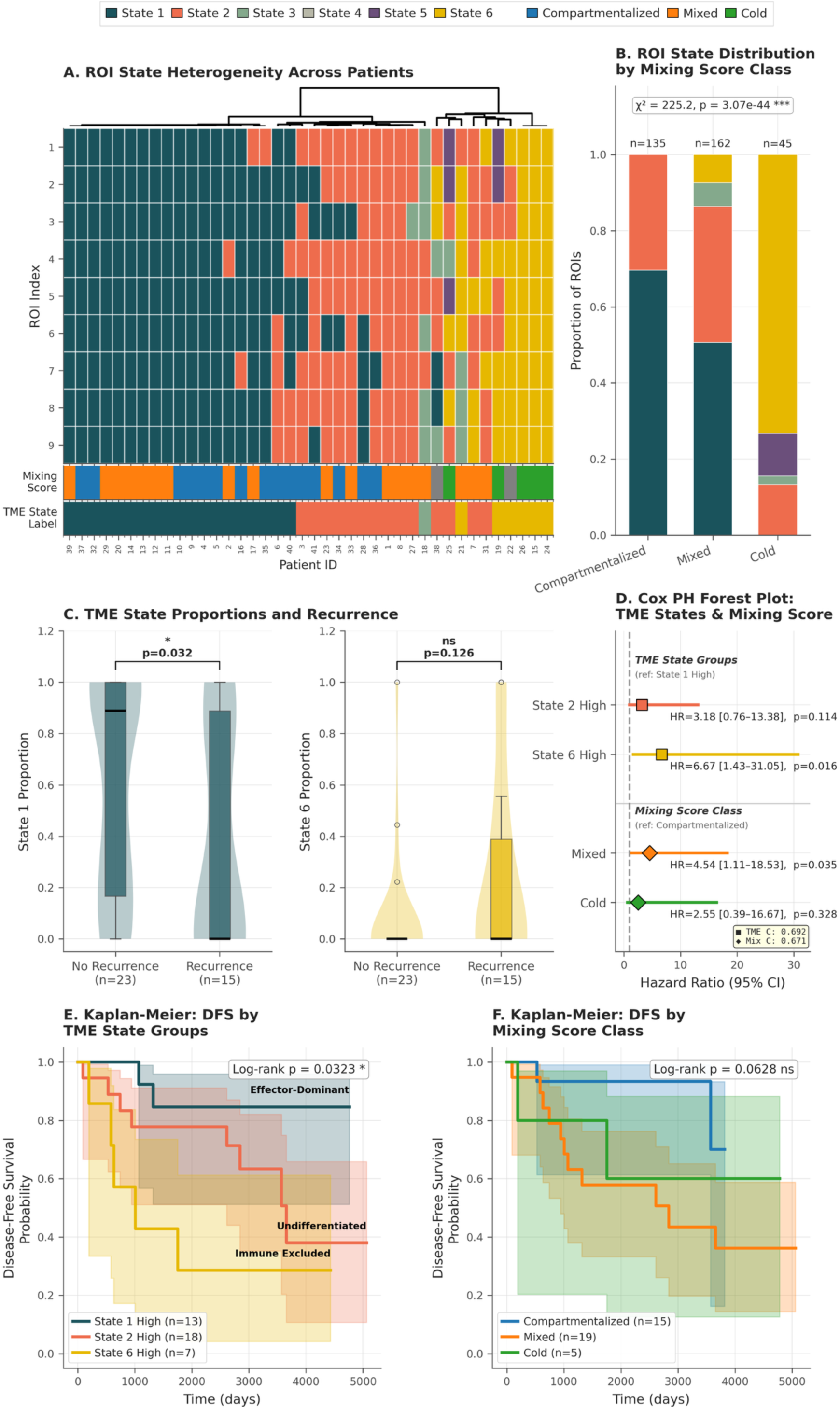
ABM-derived TME state assignments correlate with expert-derived mixing score classification and disease-free survival. **A,** TME state assignments across the MIBI cohort. Heatmap shows state assignments for ROIs (rows) across patients (columns); 9 ROIs sampled per patient. First rug plot indicates expert-derived mixing score classification (Compartmentalized, Mixed, Cold) proposed by Keren et al. Second rug plot indicates patient-level TME state assignment based on mode of ROI assignments. **B,** TME state distribution by mixing score class. Chi-square test, P < 0.001. **C,** TME state distribution by recurrence status. Higher State 1 proportions associate with absence of recurrence (P = 0.032); State 6 enrichment trends toward recurrence (P = 0.126). **D,** Multivariate Cox regression. Forest plot shows hazard ratios with State 1 as reference. State 6 shows significantly elevated hazard (HR = 7.19, 95% CI shown; likelihood ratio test P = 0.036; C-index = 0.692); Cox regression with the Mixing Score Compartmentalized category as reference. The Mixed and Cold categories indicate less elevated hazard relative to TME state (HR = 4.54 and 2.55 respectively). The model achieves a C-index of 0.671 (less explanatory), though comparison between Compartmentalized and Mixed achieves statistical significance (P = 0.035). **E,** Kaplan-Meier survival analysis. Patients with high State 1 proportions show superior disease-free survival versus those enriched in States 2 or 6 (log-rank P = 0.03). Groups with insufficient events excluded due to limited statistical power (15 total events). **F,** Kaplan-Meier survival analysis using the mixing score proposed by *Keren et al*. Patients classified as compartmentalized trend towards extended disease-free survival, however the model does not achieve statistical significance (P = 0.0628).

ABM-derived TME states for each patient significantly corresponded with mixing score classes (**Fig. 4B**; chi-square test, p < 0.001). Compartmentalized tumors were enriched for Effector-Dominant (State 1) ROIs; Mixed tumors showed intermediate compositions, and Cold tumors mapped almost exclusively to Immune-Excluded (State 6). Notably, no ROIs mapped to Exhausted-Dominant (State 4), indicating that infiltrated-but-dysfunctional TMEs are rare in this surgical resection cohort.

Despite this strong correspondence to published labels, ABM-derived states provided greater prognostic resolution. Unlike the mixing score, which assigns tumors to discrete categories, ABM states preserve continuous and within-class variation by encoding the relative prevalence of multiple biologically distinct TME configurations across ROIs. As a result, tumors sharing the same mixing score can differ substantially in their underlying ABM state composition and associated risk. Consistent with this, patients who experienced disease recurrence showed significantly lower numbers of Effector-Dominant (State 1) ROIs than those who remained disease-free (Mann-Whitney U test, p = 0.032) (**Fig. 4C**). Patients with higher proportions of ROIs in the Immune-Excluded (State 6) state tended to have more recurrence, consistent with reduced immune surveillance and tumor-dominant architecture, however this trend did not achieve statistical significance (p = 0.126).^8,37^

To quantify the association between dominant TME state and disease-free survival (DFS), we fit a Cox proportional hazards model with dominant TME state (State 1 High, State 2 High, or State 6 High) as the predictor and State 1 High as the reference category; no additional covariates were included given the cohort size (n=41). Cox proportional hazards modeling (**Fig. 4D**) confirmed this trend: relative to State 1 High patients, State 6 High patients had a 6.67-fold increased hazard of recurrence (HR = 6.67, 95% CI: 1.43–31.05, p = 0.019), while State 2 High showed a non-significant trend (HR = 3.18, 95% CI: 0.76–13.38, p = 0.122). The model achieved a concordance index of 0.692, with a significant overall likelihood ratio test (p = 0.036). A second model was trained using the mixing score labels to stratify patients with the Compartmentalized category set as reference. Patients labeled as ‘Mixed’ had a 4.54-fold increase hazard of recurrence (HR = 4.54, 95% CI: 1.11–18.53, p = 0.035) while ‘Cold’ labeled patients exhibited a 2.55-fold increase (HR = 2.25, 95% CI: 0.39–16.67, p = 0.328).

Kaplan-Meier analysis of DFS further demonstrated the prognostic value of the ABM-derived TME states. Given the cohort size (n=41;15 recurrence events), patients were grouped by dominant TME state composition to preserve statistical power. Stratification by dominant ABM TME state revealed significant differences in DFS (log-rank p = 0.032) (**Fig. 4E**). State 1 High patients (n=13) exhibited markedly prolonged DFS (> 13.7 years), whereas State 6 High patients (n=7) had substantially poorer outcomes (∼2.8 years). By contrast, the same model using all three categories of the mixing score trended towards but did not achieve statistical significance (**Fig. 4F**).

We compared these two systems via likelihood ratio test (LRT). ABM-derived state composition predicted disease free survival in the MIBI cohort better than the expert-defined mixing score (**Table 2**). The ABM TME state model achieved better discriminative ability with a higher C-index (0.692 versus 0.671; ΔC-index = +0.021) and was the only model to reach statistical significance in LRT against the null model (p=0.036).

**Table 2.**
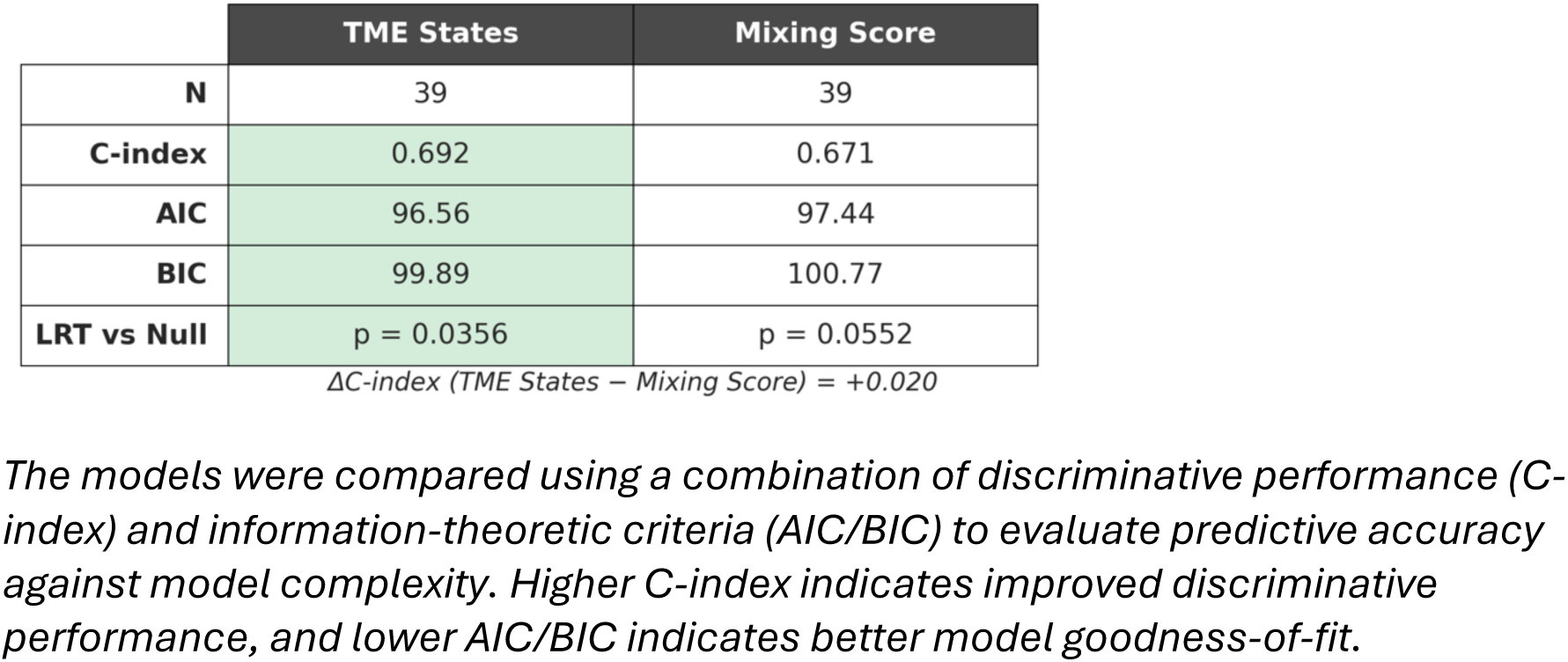
Likelihood Ratio Testing (LRT) Between TME States and Mixing Score Cox-PH Models.

### Treatment-phase TME states, rather than baseline configuration, predict immunotherapy response

Having established that ABM-derived states have the potential to capture clinically meaningful TME variation and encode potential mechanistic determinants of trajectory dynamics, we next tested whether these states predict immunotherapy response in a clinical trial setting. The NeoTRIP trial, which randomized patients to neoadjuvant chemotherapy with or without atezolizumab (anti-PD-L1), provides an ideal test case, combining longitudinal sampling with a clinically meaningful endpoint, pathologic complete response (pCR).^11,27^ We mapped 1,842 regions of interest (ROIs) from 279 patients across baseline, on-treatment, and post-treatment biospecimens to the ABM-derived TME state landscape.

Individual patients exhibited substantial spatial heterogeneity, with ROIs from the same biospecimen often mapping to multiple TME states (**Fig. 5A**). Aggregating ROI-level assignments revealed systematic shifts in dominant TME states across treatment phases: the proportion of patients classified as Effector-Dominant (State 1) increased from baseline to on-treatment before declining at post-treatment. This pattern is consistent with treatment-induced immune activation followed by either wound healing and resolution of inflammation, or exhaustion-driven divergence.^38,39^

**Figure 5.**
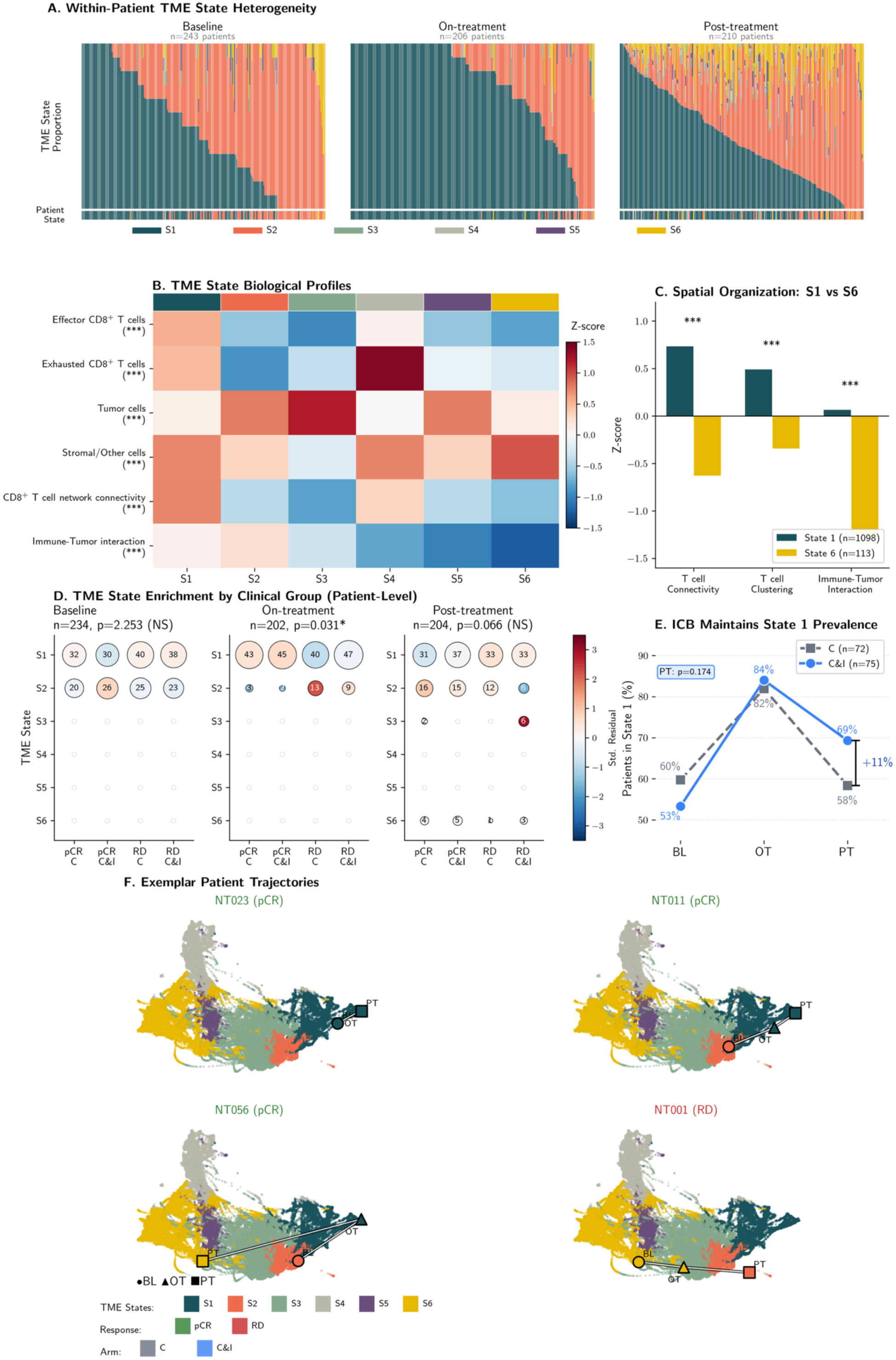
Treatment-phase TME dynamics predict immunotherapy response in triple-negative breast cancer. **A,** Within-patient TME state heterogeneity across treatment phases. Stacked bars show ROI-level state proportions per patient; rug plots indicate patient-level assignments (mode of ROI classifications). **B,** Biological validation of TME states. Heatmap shows mean z-scored features across states. Asterisks indicate Kruskal-Wallis significance after FDR correction (***P < 0.001). **C,** Spatial organization distinguishes favorable from unfavorable states. State 1 exhibits significantly higher T cell connectivity, clustering, and immune-tumor interaction frequency than State 6 (Mann-Whitney U, ***P < 0.001). n=1,098 ROIs (State 1), n=113 ROIs (State 6). **D,** Patient-level TME state enrichment by clinical group. Circle size indicates patient count; color indicates Pearson residuals. Chi-square tests show significant treatment-phase associations (on-treatment P=0.031) but not baseline (P=2.25) or post-treatment (P=0.066) after Bonferroni correction. States with <5 observations excluded to meet Chi-square assumptions. **E,** ICT maintains Effector-Dominant prevalence through treatment. C&I arm shows higher State 1 retention at post-treatment versus chemotherapy alone (69% vs. 58%; Fisher’s exact P=0.166). Complete trajectories: C n=72, C&I n=75 patients. **F,** Exemplar patient trajectories through ABM-derived TME state landscape. UMAP projections show ABM time windows (background) with clinical trajectories overlaid (circles: baseline; triangles: on-treatment; squares: post-treatment). Representative patterns include stable favorable (NT023, 1→1→1), recovery (NT011, 2→1→1), resolved inflammation (NT056, 2→1→6), and failed activation (NT001, 6→6→2). C, chemotherapy alone; C&I, chemotherapy plus atezolizumab; pCR, pathologic complete response; RD, residual disease.

To confirm that ABM-derived TME states reflect biologically meaningful TME phenotypes rather than arbitrary computational clusters, we examined cell population densities and spatial organization metrics calculated from the patient samples across all six states (**Fig. 5B**). Patient samples within each state exhibited a distinct biological profile, with highly significant differences across all features (Kruskal-Wallis p < 0.001). Clinical samples mapped to State 1 (Effector-Dominant) displayed elevated effector CD8⁺ T cell infiltration (z-score +0.54) and high T cell network connectivity (z-score +0.74), consistent with an immunologically active microenvironment. In contrast, samples mapped to State 6 (Immune-Excluded) showed depleted effector T cells (z-score −0.81) and markedly reduced immune-tumor interaction frequency (z-score −1.25). Notably, patient samples mapped to State 4 exhibited the highest exhausted T cell signature (+1.38), identifying a distinct infiltrated-but-dysfunctional phenotype rather than simple immune absence.

ABM-derived states encoded not only cell composition but also multicellular architecture. Direct comparison of spatial metrics between State 1 and State 6 confirmed profound architectural differences, with State 1 exhibiting significantly higher T cell connectivity, clustering coefficients, and immune-tumor interaction frequencies compared to State 6 (Mann-Whitney U test, p < 0.001 for all comparisons) (**Fig. 5C**). These features capture information inaccessible to biomarkers based on cell counts alone, consistent with prior work identifying spatial organization as a determinant of immunotherapy response.^10,11,15,40^

We next examined whether TME state distributions associated with treatment groups and clinical outcomes. To evaluate the relationships between these categorical variables, we applied contingency analysis with Bonferroni correction to investigate the interplay between treatment arm, patient outcome, and biospecimen phase. At baseline, analysis revealed no significant association between the predominant TME state and clinical outcome (χ² test, p_adj_ > 1.00) indicating that baseline TME configuration alone lacks sufficient predictive power for patient response (**Fig. 5D**) – consistent with prior studies.^15,25^ In contrast, significant associations between predominant TME state and treatment response emerged at the on treatment timepoint (p_adj_ = 0.031), and trended toward significance post-treatment (p_adj_ = 0.066). Patients achieving pCR showed enrichment for Effector-Dominant (State 1) classifications, while those with residual disease were enriched for Tumor-Dominant (State 2) and Immune-Excluded (State 6) states. These results support a trajectory bifurcation model in which treatment-induced TME remodeling, rather than baseline configuration, distinguishes responders from non-responders.^14^

Given that State 1 (the Effector-Dominant state) was associated with favorable outcomes, we next investigated whether immunotherapy affected the maintenance of this phenotype across treatment phases (**Fig. 5E**). Both treatment arms showed marked enrichment of State 1 at on-treatment (∼82–84% of patients), indicating that chemotherapy alone induces substantial immune activation and consistent with prior investigations into the pro-inflammatory effects of cytotoxic agents such as chemotherapy and radiation.^41,42^ However, the treatment arms diverged at post-treatment: patients receiving chemotherapy plus immunotherapy (C&I) exhibited higher persistence of the Effector-Dominant (State 1) compared to chemotherapy alone (69% of patients vs. 58%, +11%; Fisher’s exact p = 0.174). Although not statistically significant, the magnitude and direction of the effect suggest that checkpoint blockade may act primarily to sustain favorable immune states rather than initiate them. This observation supports a maintenance hypothesis for immune checkpoint therapy (ICT), in which therapeutic benefit arises from preventing transition from Effector-Dominant state induced by chemotherapy to less favorable states by slowing exhaustion accumulation, rather than from de novo immune activation.^43–46^

In addition, we observed that identical terminal TME states could be associated with divergent clinical outcomes, underscoring the limitations of static state assignment. A unique aspect of this clinical cohort is that we have MTI from repeat biospecimens, allowing us to use the integrated model and patient data to use the dynamics to resolve why similar states can have distinct outcomes. We examined individual patient trajectories through the ABM-derived TME state landscape (**Fig. 5F, Supplementary Fig. 8**) by tracing the route taken by their three biospecimens on the TME state landscape. Four archetypal patterns emerged: a stable Effector-Dominant trajectory [1→1→1] and a trajectory of improvement [2→1→1] were associated with pCR; by contrast, two patients (NT056 and NT161) terminating in Immune-Excluded (State 6) exhibited opposite clinical outcomes depending on trajectory context. Patient NT056 (pCR, C) followed a resolved inflammation trajectory [2→1→6], consistent with tumor clearance followed by wound healing and tissue remodeling. ^47–49^ Supporting this interpretation, pCR patients assigned to TME state 6 at post-treatment exhibited significantly higher fibroblast-to-effector CD8^+^ T cell ratios than patients with residual disease (**Supplementary Fig. 9**). Conversely, patient NT001 (RD, C&I) followed an immune exclusion trajectory [6→6→2], reflecting incomplete or non-response to therapy.^49–51^ These findings demonstrate that trajectory history – information invisible to single timepoint biomarkers – is essential for interpreting TME states, therapeutic response, and clinical outcomes.^43,50,52,53^

### Within-state dynamics reveal exhaustion as a mechanistic driver of trajectory divergence

Having validated that ABM-derived TME state landscape captures clinically meaningful TME variation, we next used the computational model to identify potential mechanistic drivers that govern whether a simulation follows a favorable or unfavorable tumor-immune trajectories. Sorting simulation trajectories according to similarity of their temporal progression through TME states revealed a small number of canonical transition patterns (**Fig. 6A**). Simulations progress from the Undifferentiated and the Pre-exclusion Transitional state (State 3) before progressing toward either the Pre-Exhaustion Transitional state (State 5) or toward the terminal Immune-Excluded state (State 6). Entry into the Effector-Dominant state (State 1) occurred almost exclusively from the Undifferentiated state, indicating a narrow window for effective immune activation and simulations occupying Effector-Dominant state (State 1) either remained stable or transition to the Exhaustion-Dominant state (State 4), with only a minority ultimately progressing to the Immune Excluded state.

**Figure 6.**
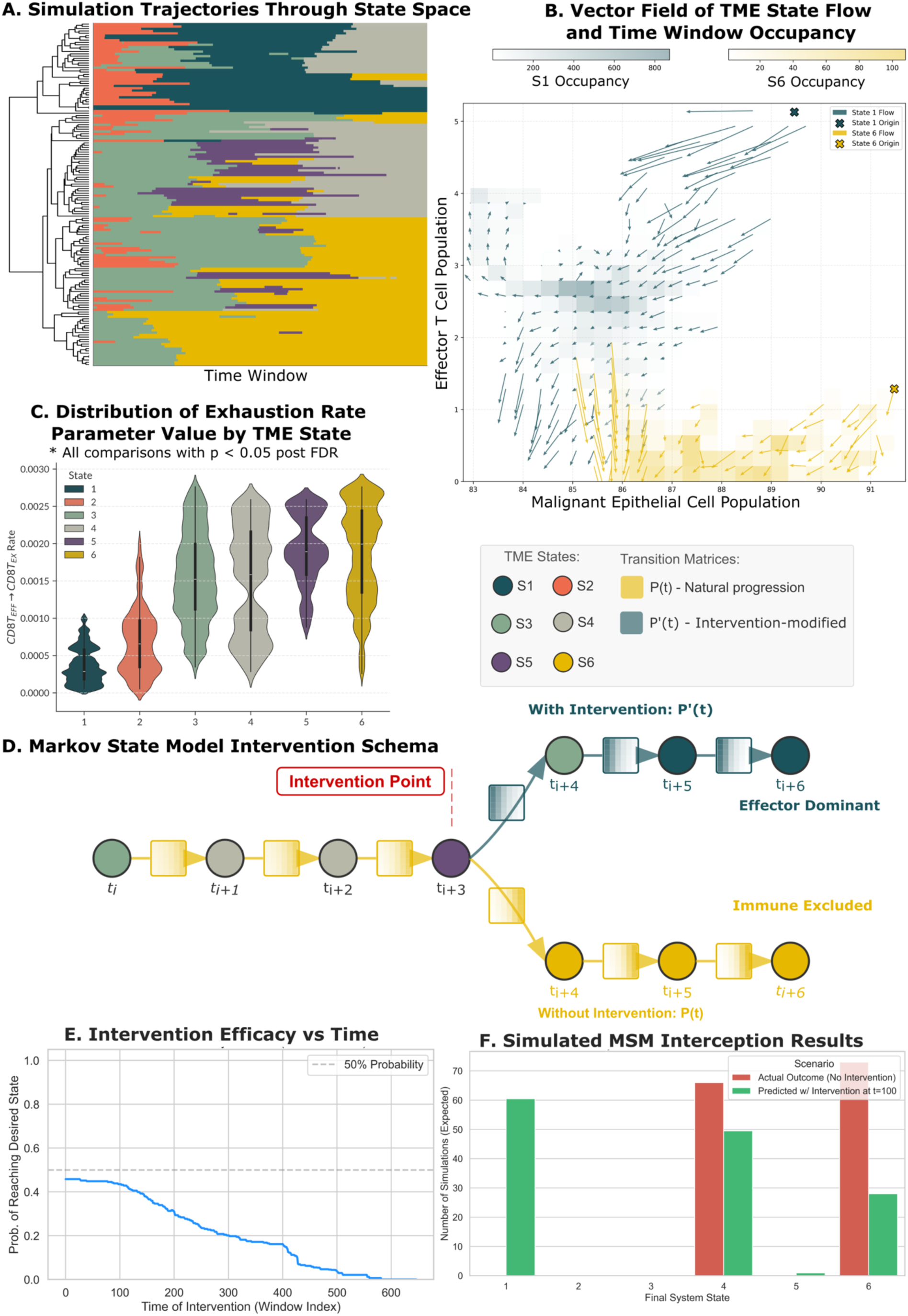
Within-state dynamics reveal mechanistic drivers of trajectory divergence and enable computational intervention modeling. **A,** Clustermap of simulation trajectories through TME states. Rows represent individual simulations; columns represent time windows. **B,** Superimposed vector fields of population-level dynamics for effector CD8^+^ T cells and malignant epithelial cells. Heatmaps show the number of time windows within each state that occupy each region of the phase space. Arrows represent the direction and magnitude of the system’s temporal evolution. Crosses indicate the initial conditions for when the first simulations enter a state, highlighting the divergent trajectories toward distinct meta-stable attractors. **C,** CD8⁺ T cell exhaustion rate distribution across states. All pairwise comparisons versus State 1 are significant (Mann-Whitney U, BH-FDR corrected, all P < 0.05). **D,** Markov state model (MSM) surrogate for intervention testing. Transition probabilities conditioned on parameter values enable rapid evaluation of treatment timing without full ABM re-simulation. **E,** Intervention efficacy versus treatment initiation time. Y-axis shows probability of reaching State 1 under simulated treatment (exhaustion rate = 0.0003). **F,** The predicted treatment outcomes using the MSM. Red bars: original terminal state distribution for simulations ending in Exhausted-Dominant (State 4) or Immune-Excluded (State 6). Green bars: predicted terminal states following simulated intervention.

To understand the mechanistic determinants governing fate along the TME State 1 – State 6 axis, we computed the vector field of population-level dynamics for simulations occupying each state, representing the direction and rate of change in effector CD8+ T cell and malignant epithelial cell abundances over each simulation time interval (**Fig. 6B**, see Methods). In this representation, arrows indicate where the tumor-immune system is heading and how fast, with shading reflecting how long simulations occupy each region. Some tumor-immune configurations act as preferred destinations to which the system is repeatedly drawn and where it tends to persist; these regions are termed meta-stable attractors. State 1 and State 6 occupy fundamentally distinct such regions with non-overlapping trajectory pathways. Simulations entering State 1 exhibit a prominent divergence: one subset consolidates around a meta-stable attractor defined by high CD8+ effector T cell infiltration and successful tumor suppression, while another bypasses this attractor, progressing toward low effector T cell density despite partial tumor control. State 6 simulations converge onto a meta-stable attractor in a high tumor-burden region with negligible effector CD8+ T cell activity, with no trajectories escaping toward immune engagement. This separation of flow indicates that State 1 and State 6 represent ingrained differences in the underlying directional forces governing the tumor-immune ecosystem, rather than transient fluctuations in cell counts.

Within TME State 1, the effector CD8^+^ T cell and tumor cell populations exhibited coupled dynamics: gradual tumor expansion triggered rapid expansion of effector CD8⁺ T cells, followed by a slower contraction phase as tumor burden decreased (**Supplementary Fig. 8A**). This mirrors predator-prey dynamics: as tumors grow, they stimulate immune expansion; as the immune response clears tumor cells, it loses its stimulus and contracts. Critically, rather than oscillating indefinitely or collapsing, these coupled dynamics damped over time and converged to a stable state in which effector T cells maintained sustained tumor control. In the model, “equilibrium” means the system reached a fixed point where cell populations stopped changing — effector T cell killing was exactly balanced by tumor cell proliferation, holding both populations steady. Strikingly, this mathematical equilibrium corresponds directly to the “equilibrium” phase of cancer immunoediting, suggesting that immune-controlled states can emerge as stable dynamical outcomes of the tumor-immune interaction rather than transient peaks.^54^

In contrast, simulations occupying State 6 showed monotonic effector CD8^+^ T cell decline toward depletion, accompanied by stabilization of tumor populations at elevated levels. The smooth and effectively irreversible nature of this decline is consistent with a tipping point dynamic or threshold beyond which immune recovery becomes unlikely without intervention.^55,56^ Spatial dynamics mirrored these population trends (**Supplementary Fig. 8B**). Effector-Dominant trajectories showed increasing effector CD8⁺-tumor interaction strength over time, alongside a progressive reduction in tumor cell degree centrality, reflecting immune-mediated fragmentation of tumor architecture. In Immune-Excluded trajectories, tumor centrality and clustering metrics remain high and stable, consistent with consolidated tumor organization in the absence of effective immune pressure. Together, these spatial signatures provide quantitative correlates of the biological processes, distinguishing durable immune control from immune escape.

To evaluate the influence of ABM input parameters on simulation trajectory fate, we examined the relationship between parameter values and TME state occupancy (**Supplementary Fig. 8C**). Consistent with other computational efforts, among all parameters, the CD8^+^ T cell exhaustion rate exhibited the highest variability across TME states.^57^ Simulations reaching Effector-Dominant (State 1) possessed low exhaustion rate input parameters, whereas simulations progressing to Late-Transitional (State 5) or Immune-Excluded (State 6) exhibited markedly elevated exhaustion rates. Within the ABM, this parameter directly controls the rate at which effector CD8⁺ T cells transition to the exhausted phenotype, highlighting a potential modifiable factor with a mechanistic role underlying the loss of immune control.

Comparing the distributions of the exhaustion rate parameter between states confirmed the input parameter value differed significantly across states (**Fig. 6C**) with all pairwise comparisons between Effector-Dominant (State 1) and other states remaining significant after Benjamini-Hochberg correction. These results associate T cell exhaustion rate with driving trajectory divergence and pose the parameter as a potential candidate for therapeutic intervention. Notably, simulations entering Immune-Excluded (State 6) also exhibited significantly lower tumor cell self-adhesion. This combination may permit immune infiltration while preventing stable immune-tumor engagement, a scenario previously associated with CD8^+^ T cell exhaustion in the absence of supportive immune niches.^8,20,58^

### Markov state modeling enables computational testing of trajectory-directed interventions

Identification of T cell exhaustion rate as a potential determinant of trajectory fate suggested that modulating this parameter – conceptually analogous to immune checkpoint blockade – could redirect unfavorable tumor-immune dynamics toward favorable outcomes. To systematically explore how the timing and magnitude of such interventions could influence trajectory outcomes, we constructed a Markov state model (MSM) as a reduced representation of the ABM-derived dynamics.^59^

The MSM provides a coarse-grained description of tumor-immune evolution by learning state-to-state transition probabilities within the simulated timeframe from the ABM trajectories. For each time window, we computed the probability of a transition between any two TME states. To simulate treatment and non-treatment regimes, we conditioned the transition probabilities on the average value of the CD8^+^ T cell exhaustion rate parameter for each TME state. We represented a simulated ICT with the between TME state transition probabilities associated with the average value of the CD8^+^ T cell exhaustion rate parameter for the Effector-Dominant TME state. (**Supplementary Fig. 11**). Reducing the progression of ABM simulations through the TME states to the simplified representation of the MSM enables rapid evaluation of potential intervention scenarios by modifying transition probabilities to reflect altered exhaustion rates, while preserving the dominant dynamical structure learned from the ABM. We verified the fidelity of the MSM to the ABM by calculating the Kullback-Leibler Divergence between the distributions of states observed in ABM output and those predicted by the MSM at each timepoint, confirming that the divergence never rose above 0.16 (average KL-divergence = 0.038, **Supplementary Fig. 12**). We then use the MSM to simulate each in silico tumor’s trajectory through the TME state landscape by predicting it’s TME state at each point in time under different treatment regimens for the duration of the simulation (**Fig. 6D**).

Using the MSM, we evaluated potential intervention efficacy as a function of treatment initiation time (**Fig. 6E**). Simulated treatment corresponding to strong checkpoint blockade (exhaustion rate = 0.00051 min⁻¹) yielded a 40–50% probability of reaching the Effector-Dominant (State 1) when initiated early. This probability declined after approximately 100 MSM intervals (where each interval represents a step along the inferred TME trajectory the length of the time-delay embedding window; see Methods), corresponding to early simulation phases, and collapsed around intervals 400–425, approaching zero by interval 550. These results define a finite therapeutic window during which trajectory interception remains feasible.

To quantify the extent to which trajectories leading to states with low effector CD8^+^ T cell presence could be redirected, we identified 139 simulations that terminated in the Exhausted-Dominant (State 4) or Immune-Excluded (State 6) TME states without intervention and simulated treatment initiation at time window 100 (**Fig. 6F**). Of these, 61 simulations (44%) were redirected to Effector-Dominant (State 1), with most rescued trajectories originating from the Immune-Excluded state (State 6) rather than from Exhausted-Dominant (State 4) configurations. Together, these results demonstrate the potential of a reduced, ABM-derived state-space representation to enable systematic exploration of intervention timing and efficacy, without requiring full ABM re-simulation.

## Discussion

In this study, we present a trajectory-centric computational framework for interpreting static tissue imaging data through the lens of mechanistic tumor-immune dynamics. We use an ABM of the tumor-immune interaction in the TME and apply naïve exploration of ABM parameter space to learn a mechanistically constrained reference landscape of TME dynamics. We investigate this landscape by evaluating how experimentally quantified clinical tumor tissue samples, collected via regular biopsy and post-surgical pathology, distribute across its varied regions.

Our key methodological advance is the use of ABM ensemble simulation to generate the reference dynamical landscape – a coordinate system that exists independent of any specific patient or cohort, and onto which new clinical samples can be projected without re-simulation. While we demonstrate this framework in TNBC using PhysiCell-based rules, the approach is generalizable to any cancer type for which an ABM encoding relevant cell populations and interactions is available; the parameter exploration, state identification, and clinical projection steps are universal.

A central finding of this study is that TME states assessed during or after treatment outperform baseline configurations in predicting clinical response. In the NeoTRIP cohort, baseline TME states showed no significant association with pathologic complete response, whereas on-treatment states robustly distinguished responders from non-responders. This pattern supports a trajectory bifurcation model: baseline states represent pre-perturbation conditions with multiple possible fates, while treatment-phase states encode the system’s committed response to therapy. These results suggest that early on-treatment biospecimens may provide more actionable biomarkers for adaptive treatment strategies than pre-treatment assessment alone.^11,60^

Our findings further suggest that immune checkpoint blockade in TNBC acts primarily by maintaining favorable immune configurations that were associated with chemotherapy effects rather than initiating immune activation de novo. Both treatment arms, chemotherapy alone vs chemotherapy plus ICT, exhibited strong induction of Effector-Dominant states on-treatment (i.e. ∼14 days after starting therapy), indicating that chemotherapy alone is sufficient to drive substantial immune activation. However, checkpoint blockade was associated with greater persistence of this phenotype at post-treatment. This observation aligns with clinical trial data showing that checkpoint inhibitors improve long-term outcomes even in patients who do not achieve pCR,^61,62^ and supports a model in which ICT slows exhaustion-driven transitions away from immune-active states by preserving effector functionality ^44,63^. Notably, ABM parameter analysis identified T cell exhaustion rate as a dominant association with simulated trajectory fate, providing a potential mechanistic basis for this interpretation.

Critically, trajectory context proved essential for interpreting terminal TME states. Identical post-treatment Immune-Excluded configurations were associated with opposite clinical outcomes depending on prior trajectory: patients following a [2→1→6] path achieved pCR, consistent with resolved inflammation following tumor clearance, whereas patients following a [1→2→6] path had residual disease, reflecting progressive immune exclusion. This distinction is invisible to static biomarkers and demonstrates that the biological meaning of a spatial phenotype depends on its dynamical history. It follows that two serial tissue samples substantially improve the interpretability of spatial biomarkers compared to single-timepoint profiling, a finding supported by other recent findings.^12^ By reconstructing likely trajectories from the ABM-derived TME state landscape, our framework resolves this ambiguity and provides a principled basis for interpreting treatment response.

### Limitations

Several limitations warrant consideration. First, our ABM simulations span only two weeks of biological time, which is substantially shorter than months-long clinical treatment courses. As a result, correspondence between simulation time and clinical timescales remains approximate, and longer-term processes including immune memory and tertiary lymphoid structure formation are not captured. Second, the ABM necessarily simplifies biological complexity. For example, macrophage polarization is represented as a discrete axis rather than a continuum, and additional populations, including B cells, dendritic cells, and fibroblast subtypes, are not explicitly modeled. These simplifications limit the ability of the framework to identify a broader range of therapeutic targets for microenvironmental manipulation. Finally, pathologic complete response (pCR) was used as the primary clinical endpoint following NeoTRIP trial design.^27^ However, pCR is an imperfect surrogate for long-term survival ^61,62^, and validation against event-free survival and overall survival endpoints will be essential for clinical translation.

### Future Directions

The framework described herein provides a foundation for interpreting tissue biospecimens as snapshots positioned along dynamic tumor microenvironment trajectories,^14^ with potential implications for biomarker development and temporally adaptive treatment strategies. ^13^ More broadly, ensemble-simulation approaches of this kind offer a principled path toward the long-term goal of therapeutic microenvironment manipulation. By systematically exploring model parameter space, one may identify antitumor features of the TME, link those features to candidate control mechanisms, and ultimately inform strategies to increase their prevalence or functional efficacy. In the present study, this framework identified T cell exhaustion rate as the dominant mechanistic correlate of trajectory fate. This finding was consistent with the available clinical trial evidence on PD-1 checkpoint inhibition and provided a concrete, if simplified, proof of concept. At the same time, we acknowledge that the current model remains an early demonstration with limited direct clinical applicability. Future work incorporating additional cell populations such as stromal subtypes, dendritic cells, and B cells, would expand the modeled parameter space and may reveal additional targets for microenvironmental manipulation beyond T cell functional state, further extending the translational potential of this approach. Eventually, the underlying principles of this framework may guide adaptive sampling protocols, prioritizing early on-treatment tissue collection in patients with heterogeneous and rapidly evolving TMEs – a hallmark of TNBC – to resolve inherent state ambiguity.

## Resource Availability

### Lead contact

Requests for further information and resources should be directed to and will be fulfilled by the lead contact, Dr. Young Hwan Chang.

### Materials availability

This study did not generate new unique reagents.

### Data and code availability

- All data generated in this study are available within the article and available on Zenodo: (link made available after publication).
- All original data and code have been deposited at Zenodo at https://doi.org/10.5281/zenodo.19226009 and is publicly available as of the date of publication.
- The software code together with detailed instructions can be found on Zenodo: (link made available after publication), and GitHub: (link made available after publication)
- Any additional information needed to reanalyze the data reported in this paper is available from the lead author by request.

## Acknowledgments

E.M.C was supported by the NIH/NCI 1T32CA254888 and by the NIH/NIGMS 5T32GM141938. L.M.H, Y.H.C, R.H, H.L.R, and P.M were supported by the Jayne Koskinas Ted Giovanis Foundation for Health and Policy. E.J.F. was supported by the National Foundation for Cancer Research (NFCR) and E.J.F, D.R.B, R.H, H.L.R, and P.M NIH/NCI U24CA284156. GBM is supported by a kind gift from the Miriam and Sheldon Adelson Medical Research Foundation, the Breast Cancer Research Foundation and NIH UO1 CA281902. The research reported in this publication used computational infrastructure supported by the Office of Research Infrastructure Programs, Office of the Director, of the National Institutes of Health under award number S10OD034224. The content is solely the responsibility of the authors and does not necessarily represent the official views of the National Institutes of Health.

## Author Contributions

E.M.C.: conceptualization, data curation, formal analysis, investigation, methodology, project administration, software, validation, visualization, writing – original draft, and writing – review and editing. R.H.: methodology, supervision. H.L.R.: methodology, supervision. D.R.B.: supervision. J.W.G.: conceptualization, supervision, investigation, writing – review & editing. G.B.M.: supervision, methodology, funding acquisition, investigation, writing – review & editing. E.J.F.: investigation, funding acquisition, supervision, and writing – review and editing. P.M..: methodology, investigation, funding acquisition, supervision, and writing – review and editing. L.M.H.: conceptualization, funding acquisition, supervision, project administration, and writing – review and editing. Y.H.C.: conceptualization, methodology, funding acquisition, resources, supervision, project administration, and writing – review and editing.

## Methods

### Agent-Based Model

We adapted a previously published PhysiCell-based agent-based model of tumor-immune interactions to use for triple-negative breast cancer.^19,22^ The model explicitly represents six cell types: tumor cells, naïve, effector, and exhausted CD8⁺ T cells, and macrophages spanning the M0→M1→M2 polarization axis. Cell-level behaviors including proliferation, apoptosis, migration, and phenotypic state transitions are governed by mechanistic rules that encode established tumor-immune feedback processes.

Specifically, M0 macrophages differentiate to the pro-inflammatory M1 phenotype upon contact with tumor cells or apoptotic debris, subsequently secreting pro-inflammatory cytokines, including IFN-γ, to activate naïve T cells into tumor-killing effector cells. M1 macrophages can polarize to the immunosuppressive M2 phenotype, which secretes IL-10 to drive effector T cell exhaustion. This mechanistic framework captures the essential feedback loops that determine whether the immune system achieves tumor control or escapes immune surveillance.

Simulations were conducted over two weeks of biological time (336 hours), with an initial population of approximately, 1,100 cells distributed within a two-dimensional spatial domain of 2000×2000 µm. To reflect experimental temporal sampling strategies, simulation output was saved every 10 minutes of simulation time during the first 72 hours, capturing early immune-tumor dynamics, and subsequently saved at one-hour intervals for the remainder of the simulation period.^64,65^

### Parameter Sampling

Model parameters governing immune cell behavior were sampled using Latin Hypercube Sampling to enable systematic exploration of the tumor-immune response landscape (Supplementary Table 1).^66–68^ A total of fifty unique parameter combinations were generated. For each parameter set, simulations were initialized from one of three randomized initial conditions (cell positions), with configurations constrained to maintain comparable spatial relationships among cell populations (**Supplementary Figure 1A-C**).

### Spatial Statistics Calculation

At each simulation timepoint, we computed spatial statistics characterizing cell-cell interactions and population compositions using the Squidpy Python package with custom wrappers.^69^ Computed metrics included cell type proportions, degree centrality, clustering coefficients, and pairwise cell-cell interaction frequencies. Spatial cell graphs were constructed using Delaunay triangulation as implemented in Squidpy.

### Feature Normalization

Spatial features were normalized using a within-timestep procedure designed to preserve temporal dynamics while enabling comparison across independent-simulations. Specifically,

1. Features were grouped by simulation timestep.
2. A Yeo-Johnson power transformation was applied to reduce distributional skewness.
3. Z-score standardization was performed within each timestep group.
4. Original timestep means were added back to preserve the temporal signal.

This normalization procedure ensures feature comparability across simulations while retaining biologically meaningful temporal trends in spatial organization.

### Time-Delay Embedding

We applied time-delay embedding based on Takens’ theorem to transform multivariate spatial statistics time series into a high-dimensional phase space.^29,30,32^ For each simulation, consecutive timepoints were concatenated into overlapping windows of length *W* = 50 and stride *t* = 1. Each window corresponds to a single point in the embedded space, capturing the system’s recent temporal history. This embedding preserves the topological structure of the underlying dynamical system, enabling identification of recurrent dynamical configurations.

### State Identification via Clustering

We constructed a k-nearest neighbor graph over the embedded time windows and applied Leiden community detection to partition the dynamics into fine-grained clusters. These communities were then aggregated via hierarchical clustering with Euclidean distance to yield six interpretable metastable states. The number of metastable states was determined using the elbow method applied to within-cluster sum of squares (WSS), balancing interpretability with dynamical resolution (**Supplementary Fig. 2**). For visualization, Uniform Manifold Approximation and Projection (UMAP) was applied using default minimum embedding distance, with the number of nearest neighbors set to the total number of time windows per simulation (n_neighbors = 646).

### Clinical Cohort Processing

#### MIBI-TOF Cohort (Keren et al., 2018)

Single-cell segmentation data from 41 TNBC patients were obtained from the published dataset.^8^ Cell phenotypes were mapped to functional categories to corresponding to ABM cell types: CD8 T cells, macrophages, tumor cells (keratin-positive), and other category comprising remaining cell populations (**Supplementary Fig. 10**). Spatial coordinates were extracted from segmentation masks by calculating cell centroids.

Processed data were stored as AnnData objects containing marker expression, cell phenotype annotations, and spatial coordinates. The therapeutic status of these patients is not known, making these samples represent an under-contextualized but true snapshot of TME dynamics during a patient’s treatment.

#### NeoTRIP IMC Cohort (Wang et al., 2023)

Single-cell imaging mass cytometry (IMC) data from 279 patients enrolled in the NeoTRIP clinical trial patients were processed using a similar workflow.^11,27^ The publicly available imaging mass cytometry and associated clinical data were accessed from the Zenodo repository (https://doi.org/10.5281/zenodo.7990870).^70^ Cell phenotypes were harmonized to match ABM categories as follows: CD8⁺T, CD8⁺GZMB⁺T, and CD8⁺TCF1⁺T populations were mapped to "effector_T_cell"; CD8⁺PD1⁺T_Ex cells were mapped to "exhausted_T_cell"; epithelial cells were mapped to "malignant_epithelial_cell"; and all remaining cells were labeled as "other." Data were stored as AnnData objects with marker expression, phenotype annotations, and spatial coordinates.

Of note, post-treatment tissue was obtained during surgical excision after completing the neoadjuvant regimen, approximately 24 weeks from treatment initiation (after 8 3-week cycles). However, 25% of patients in each arm discontinued neoadjuvant therapy early due to adverse events (median 6 cycles; range 1–7), meaning the actual time from treatment start to surgical biopsy varied across patients. Patient-level timing data are not reported in the primary trial publications.^11,27^

### ROI Sampling Strategy

To address the Modifiable Areal Unit Problem arising from differences in field-of-view dimensions between clinical datasets and ABM simulations, we developed a standardized region-of-interest (ROI) sampling strategy.^34,35^ As the FOV sizes and geometries varied across data sets and collection timepoints, dataset-specific but conceptually aligned sampling procedures were applied.

For the Keren/MIBI dataset, we generated candidate ROI centers using spatial Poisson sampling, which constructs a grid over the sampling region and selects an ROI center per grid space. ROIs were retained only if the intersection-over-union (IoU) with all previously selected ROIs was below a defined threshold (IoU < 0.5), computed using pairwise comparison. This approach effectively tiles each FOV while limiting excessive overlap, capturing local spatial heterogeneity that would otherwise be averaged out in whole-slide analysis.^35,71^

As the MIBI/Keren data set maintained consistent FOV dimensions across all patient samples, ROI center locations were determined using the first sample and subsequently applied to all remaining samples. ROI sizes between 200 µm and 2000 µm were tested to identify dimensions that best matched the number of cells within each ROI to the number of cells initialized in the ABM initialization. This optimization yielded an ROI dimension of 800 x 800 µm, resulting in nine ROIs per patient sample (**Supplementary Fig. 6**, **Fig. 3B**).

In contrast, image acquisition methods in the NeoTRIP (Wang *et al.*) data set varied substantially across timepoints and patients, spanning surgical resections to fine needle biopsies.^27^ As a result, both the shape and size of sampled tissue regions differed across samples (e.g., multiple biopsies collected from a single patient at the same time point could have distinct dimensions). We therefore applied the same ROI size optimization procedure used for the MIBI dataset, identifying at ROI dimensions of 300 x 300 pixels that maximized (1) the number of biopsies eligible for ROI extraction, and (2) the match of the average number of cells within each ROI to the number of cells present within the ABM initial conditions.

### Clinical ROI Spatial Analysis

For each sampled ROI, spatial cell-cell graphs were constructed using Delaunay triangulation. Spatial statistics were then computed using the same pipeline applied to ABM simulations. Features were normalized using the same within-group procedure, consisting of Yeo-Johnson power transformation, Z-score standardization, and re-centering with original timestep means.

### State Landscape Mapping

Clinical ROIs were projected into the ABM-derived TME state landscape using k-nearest neighbor classification (k=1) with cosine distance. Macrophage-associated features were excluded from the mapping procedure, as neither clinical antibody panel included markers enabling discrimination of M1 and M2 polarization states.

To validate this mapping strategy, 15% of ABM time windows were held out, macrophage features were removed, and the remaining windows were classified using the same k-nearest neighbor framework. Ground truth labels for each time step were obtained by taking the mode of the TME state labels for each window it belonged to in the time delay embedding. We then applied the mapping strategy to each held-out time step and assigned it the corresponding TME state label. Mapping accuracy was assessed by agreement with the ground truth TME state labels, yielding 97% concordance (**Supplementary Fig. 3**). Patient-level state assignments were determined by the mode of TME state labels across constituent ROIs. For analyses requiring a single state per patient-timepoint (e.g. multiple biospecimens collected at a single time point), the mode of ROI classifications was used.

### Vector Field and Occupancy Analysis for TME States 1 and 6

To characterize the underlying dynamical manifolds and flow of the system with regard to the effector CD8^+^ T cell and tumor cell populations, we constructed state-specific vector fields in the phase space defined by this pair of features. We first filtered for simulations that occupied TME State 1 or TME 6 at any point in their trajectories. For each simulation trajectory, local displacement vectors (Δ*X*, Δ*Y*) were calculated between consecutive time-delayed windows. The phase space was discretized into a regular grid, and displacement vectors were aggregated based on their corresponding TME state assignment. Within each grid cell, the mean displacement vector was computed by averaging all observed (Δ*X*, Δ*Y*) values, providing an estimate of the local system velocity and flow direction. To visualize the residence time of the system within different regions of the phase space, we calculated the time step occupancy as a two-dimensional histogram of all observed state-specific time points. This combined visualization reveals the attractors, divergent trajectories, and meta-stable regions that define the distinct dynamical regimes of the simulated tumor microenvironment. Visualizations were produced using the numpy and matplotlib python packages.^72,73^

### Markov State Modeling

To enable rapid testing of intervention strategies without full ABM re-simulation, we constructed Markov state models (MSMs) from observed dynamical trajectories. At each timestep, state transition probability matrices were estimated by counting state-to-state transitions across all simulations and normalizing counts by row sums. We verify the fidelity of the MSM to the observed trajectories of full ABM simulations across the TME state landscape by calculating the Kullback-Leibler Divergence (KL Divergence), a measure of the difference between two distributions, at each time interval.

To incorporate parameter dependence, transition probabilities conditioned on model parameter values (specifically, exhaustion rate) were computed by constructing dynamics matrices from subsets of simulations falling within predefined parameter ranges at each transition interval.

Therapeutic intervention was simulated by modifying transition probabilities to reflect reduced exhaustion rates corresponding to strong immune checkpoint blockade effect (0.016 min⁻¹), using the parameter-matched dynamics matrix for each transition.

Intervention efficacy was evaluated by propagating initial state distributions through the modified transition matrices and computing the resulting probability of reaching the Effector-Dominant state.

## Statistical Analysis

Associations between categorical variables, including TME state, clinical outcome, and treatment arm, were assessed using chi-square tests with Pearson standardized residuals. Comparisons of continuous variables were performed using Mann-Whitney U tests. Disease-free survival was analyzed using Kaplan-Meier estimators with log-rank tests and Cox proportional hazards regression. Model performance comparisons were evaluated using the concordance index (C-index), Akaike Information Criterion (AIC), and Bayesian Information Criterion (BIC). Multiple hypothesis testing was corrected using the Benjamini-Hochberg false discovery rate procedure. All statistical analyses were performed in Python using scipy, lifelines, and statsmodels packages.^74–76^

## Data and Code Availability

Mibi images and clinical data from Keren et al. may be accessed from https://www.angelolab.com/mibi-data and the supplemental data included in the publication.^8^ Data from Wang et al. are publicly available from the Zenodo repository associated with the publication (https://zenodo.org/records/7990870). ABM simulation code, analysis pipelines, and processed data will be available on GitHub at https://github.com/emcramer/tme-trajectory-landscape and Zenodo at https://doi.org/10.5281/zenodo.19226009 upon publication.

