## Supplemental Materials for "Tumor-immune trajectory context connects static tissue architecture to clinical outcomes" (from another version of this preprint: version 1, 2026-03-30)

### Supplementary Materials

#### Supplementary Figure 1

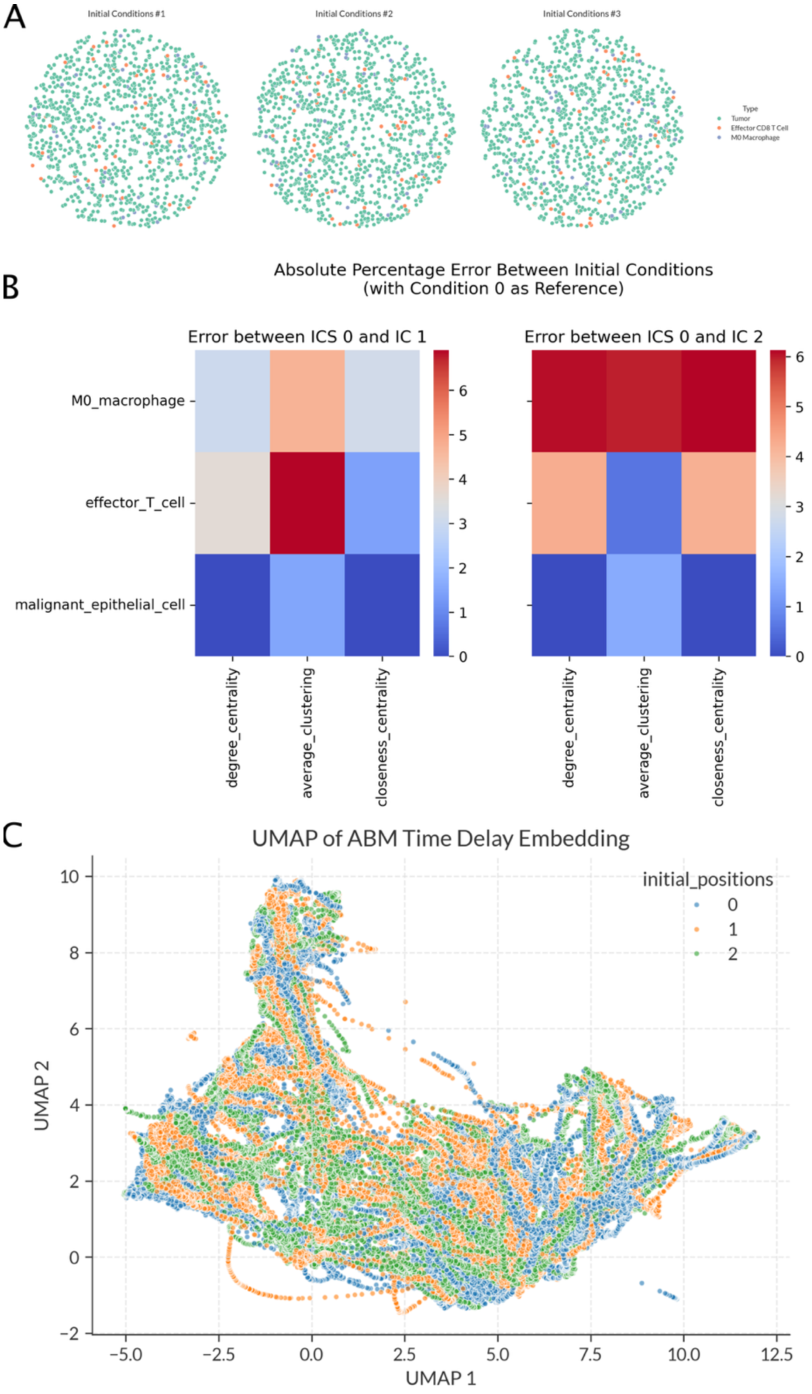

**Supplementary Figure 1. ABM Initial conditions and spatial constraints. a,** Cell positions and cell phenotypes used to initialize the ABM simulation. **b,** Centrality scores were used to control initial conditions for similarity. Heatmaps show absolute error between centrality scores. Error threshold for initial cell positions was set to 10%. **c,** UMAP projection of the ABM state space, colored by the initial cell positions (ICs) used at the start of each simulation. No observable pattern emerges, demonstrating that ICs were not primary drivers of simulation trajectories.

#### Supplementary Figure 2

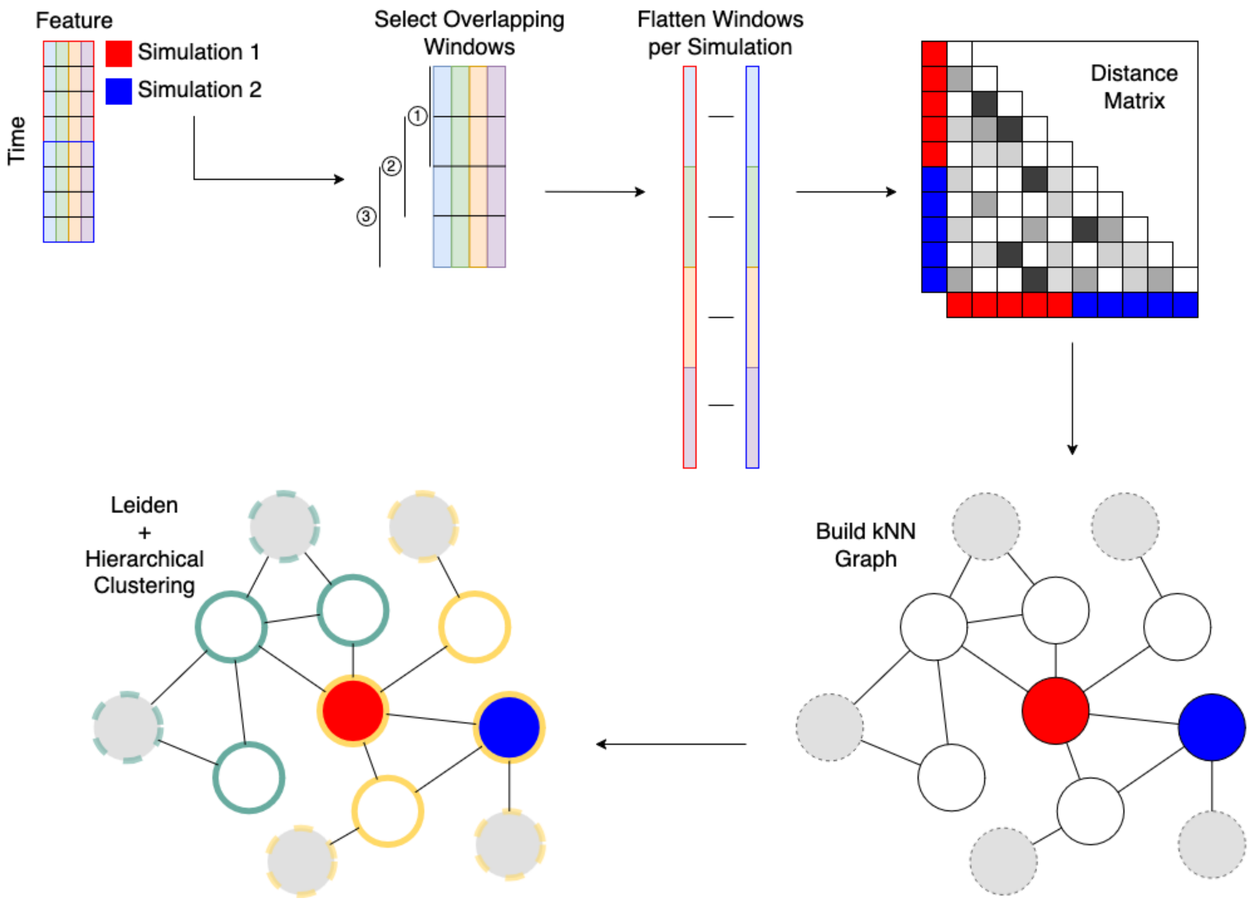


**Supplementary Figure 2. Schematic diagram of time-delay embedding and community detection/clustering approach for identifying TME states within the landscape.**

### Supplementary Figure 3

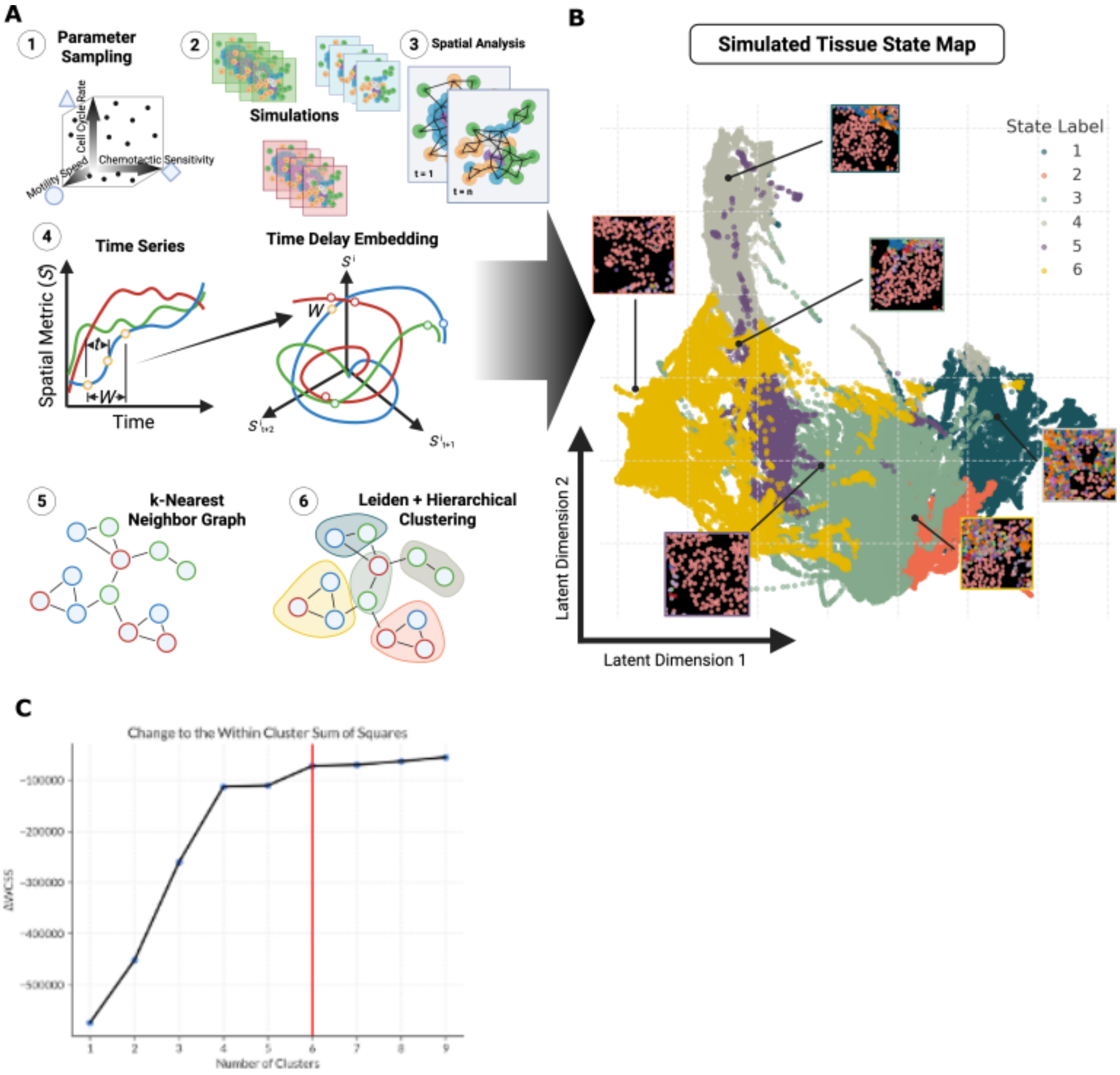


**Supplementary Figure 3. a,** Overview of the computational pipeline/workflow used to develop the TME state space landscape (see methods for full description). **b,** Resulting TME state space landscape visualized in two dimensions with UMAP and colored according to state assignment. Each dot in the UMAP plot represents a time window from a given simulation. **c,** Within Cluster Sum of Squares (WCSS) plot for hierarchical clustering**.** Empiric determination of the number of clusters to use with hierarchical clustering after Leiden community detection. The elbow method was applied using the delta of the WCSS to determine an optimal number of clusters.

### Supplementary Figure 4

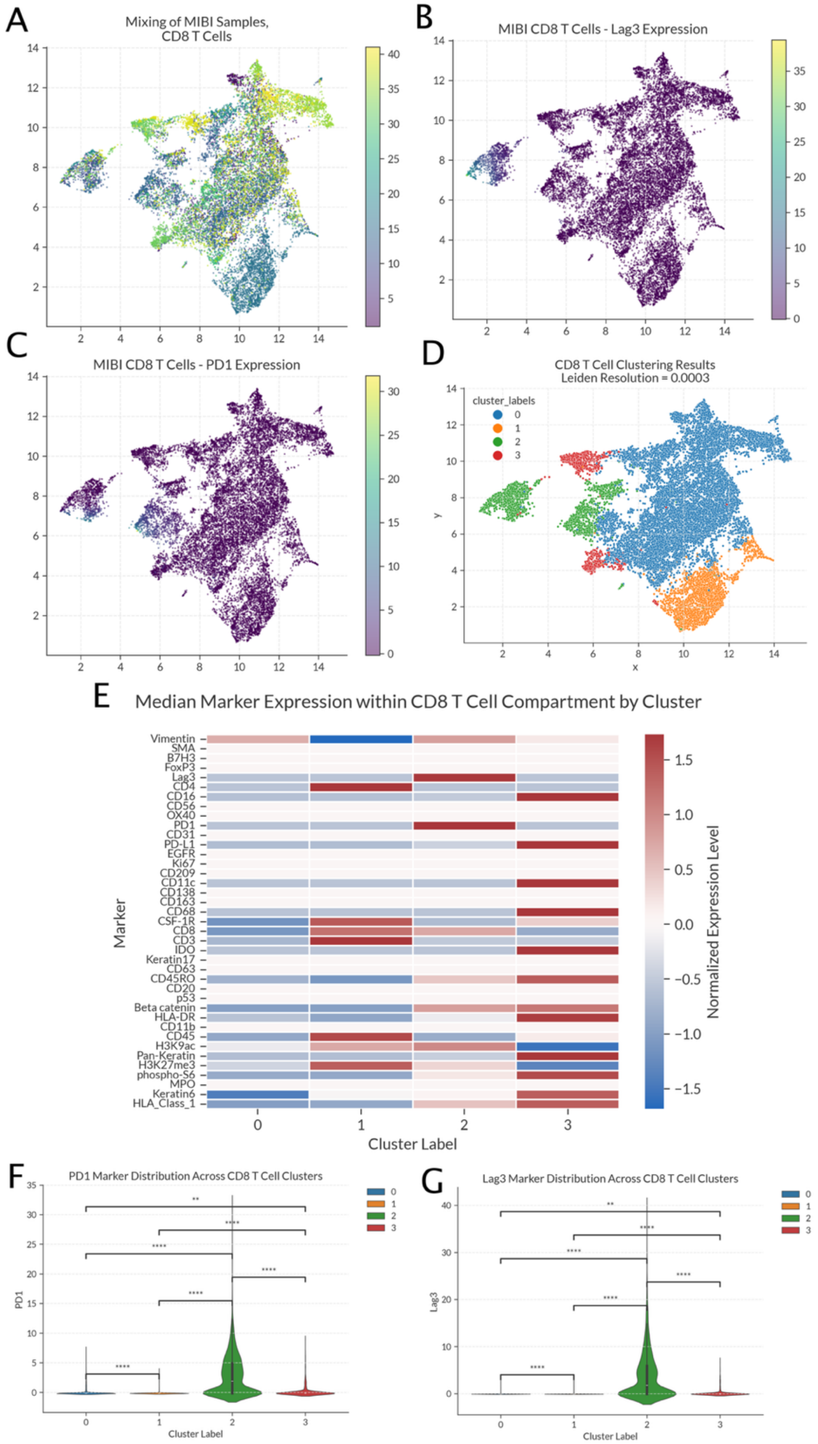


**Supplementary Figure 4. MIBI/Keren Data Set Exhausted CD8+ T Cell Functional Status Phenotyping. a,** Visualization of batch effect from each patient on the CD8+ T cell population only. Keren et al previously addressed background and batch effects before making the data publicly available. **b,** Lag3 marker expression. **c,** PD1 marker expression. **d,** Clustering results with Leiden clustering algorithm. Resolution parameter set to 0.0003. Cluster 2 appears to capture the CD8+ T cells with higher Lag3 and PD1 expression. **e,** Heatmap showing the expression levels of each marker measured with MIBI-TOF across the identified clusters. **f,** Violin plot showing the distribution of PD1 expression across the clusters. Cluster 2 had higher expression that reached statistical significance ($p<0.05$, Mann-Whitney U test). **g,** Violin plots showing the distribution of Lag3 expression across the clusters. Cluster 2 had higher expression that reached statistical significance ($p<0.05$, Mann-Whitney U test).

### Supplementary Figure 5

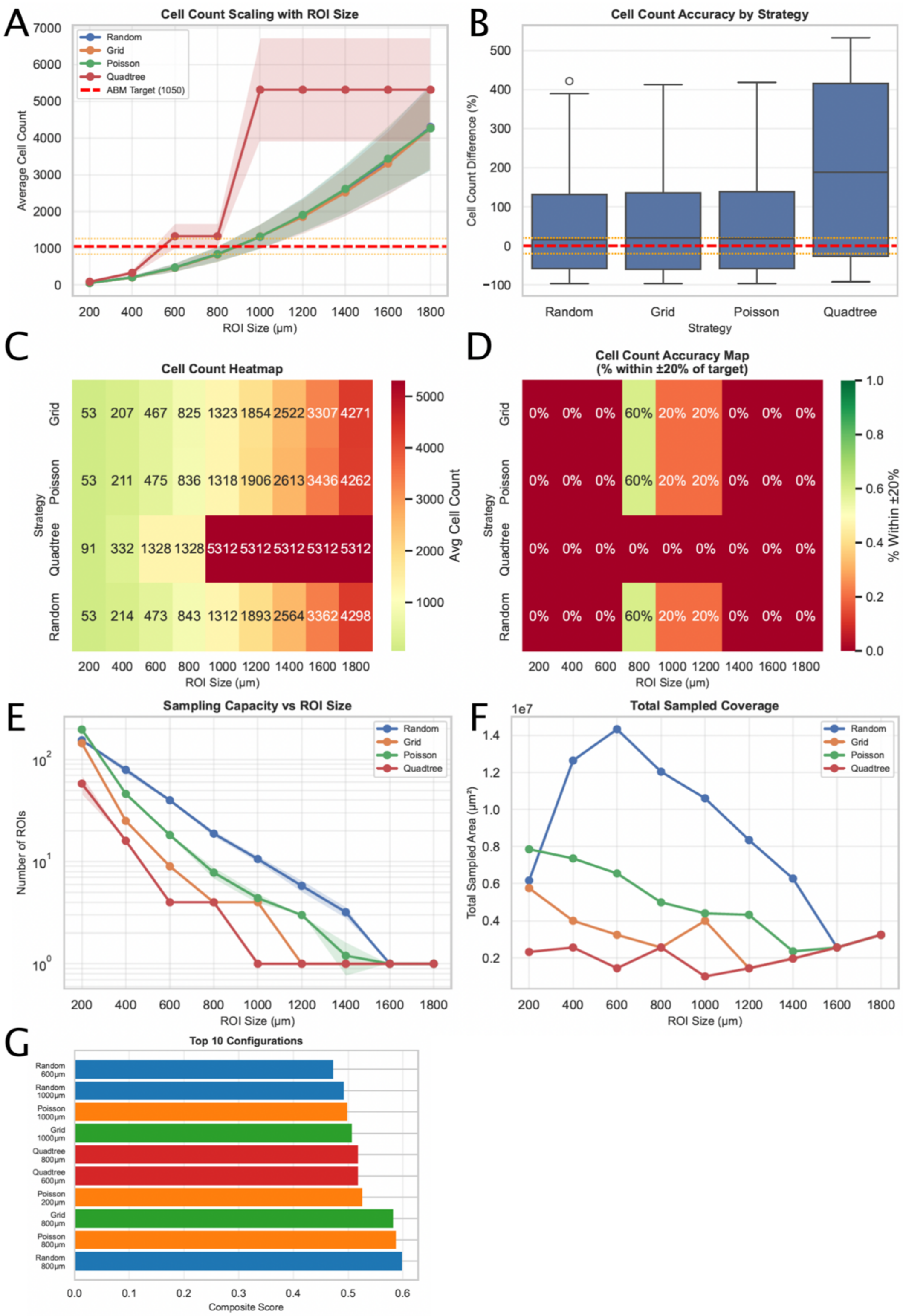


**Supplementary Figure 5. Optimization for ROI Selection Strategy on MIBI-TOF/Keren Dataset.**

#### Supplementary Figure 6

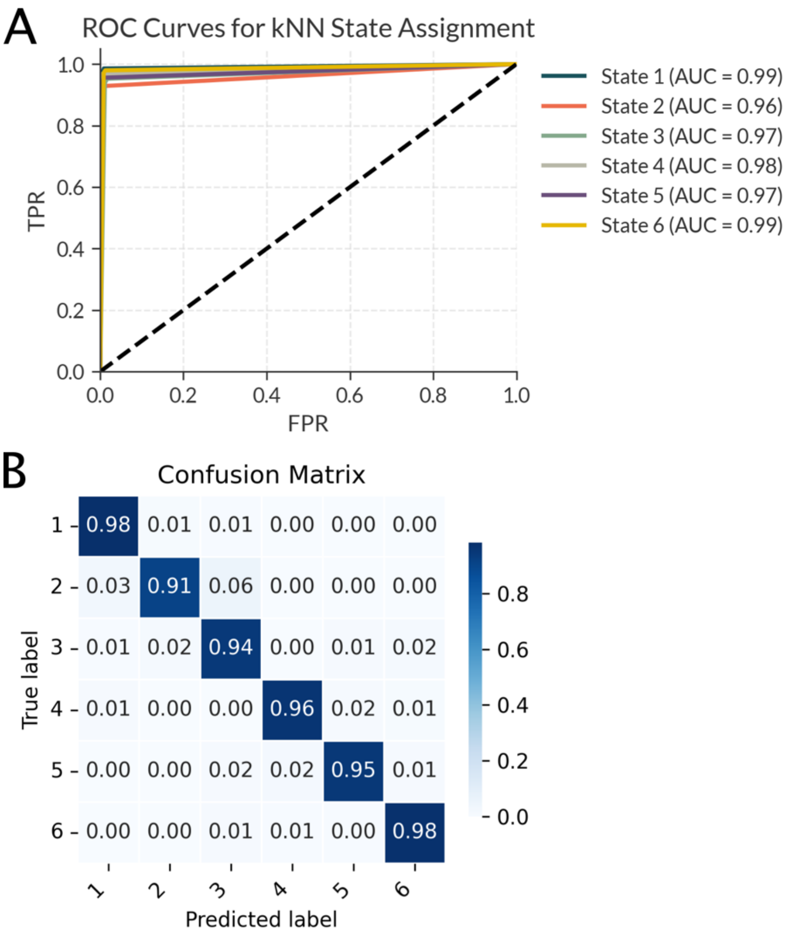


| True label | 1 | 2 | 3 | 4 | 5 | 6 |
|---|---|---|---|---|---|---|
| 1 | 0.98 | 0.01 | 0.01 | 0.00 | 0.00 | 0.00 |
| 2 | 0.03 | 0.91 | 0.06 | 0.00 | 0.00 | 0.00 |
| 3 | 0.01 | 0.02 | 0.94 | 0.00 | 0.01 | 0.02 |
| 4 | 0.01 | 0.00 | 0.00 | 0.96 | 0.02 | 0.01 |
| 5 | 0.00 | 0.00 | 0.02 | 0.02 | 0.95 | 0.01 |
| 6 | 0.00 | 0.00 | 0.01 | 0.01 | 0.00 | 0.98 |

**Supplementary Figure 6. Mapping test with modified ABM output. a,** Receiver Operating Characteristic (ROC) curves of the kNN mapping method. 15% of the ABM time steps were randomly sampled, had macrophage-related information removed from their vectors, and their TME state labels used as ground truthy to test mapping prediction. kNN with k=1 was used to map modified time step windows back to their source windows, and the TME state labels of the assigned windows were compared to ground truth labels. **b,** Confusion matrix for the kNN classifier used for the mapping procedure. Prediction accuracy exceeded 90% for each TME state.

### Supplementary Figure 7

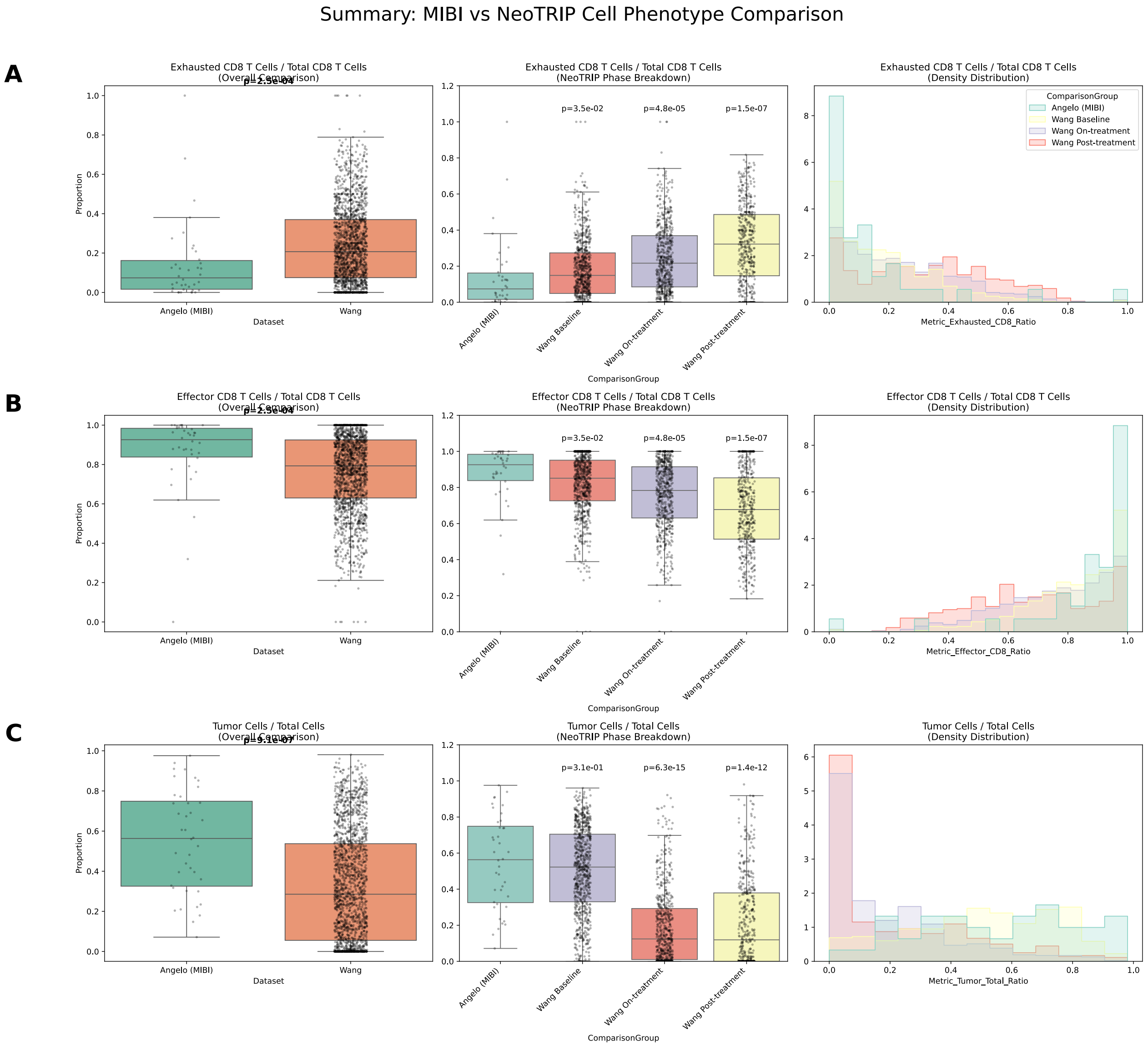


**Supplementary Figure 7. Key Cell Type Differences Between MIBI-TOF and IMC Datasets. a,** Exhausted CD8+ T cell relative population comparison. Left: comparison of distributions in each ROI by Kruskall-Wallis test. Middle: breakdown comparing populations across NeoTrip biopsy phases. Right: visualization of distributions. **b,** Effector CD8+ T cell relative population comparison. Identical Left/Right pattern as panel A. **c,** Tumor cell relative population comparison. Identical Left/Right pattern as panel A.

### Supplementary Figure 8

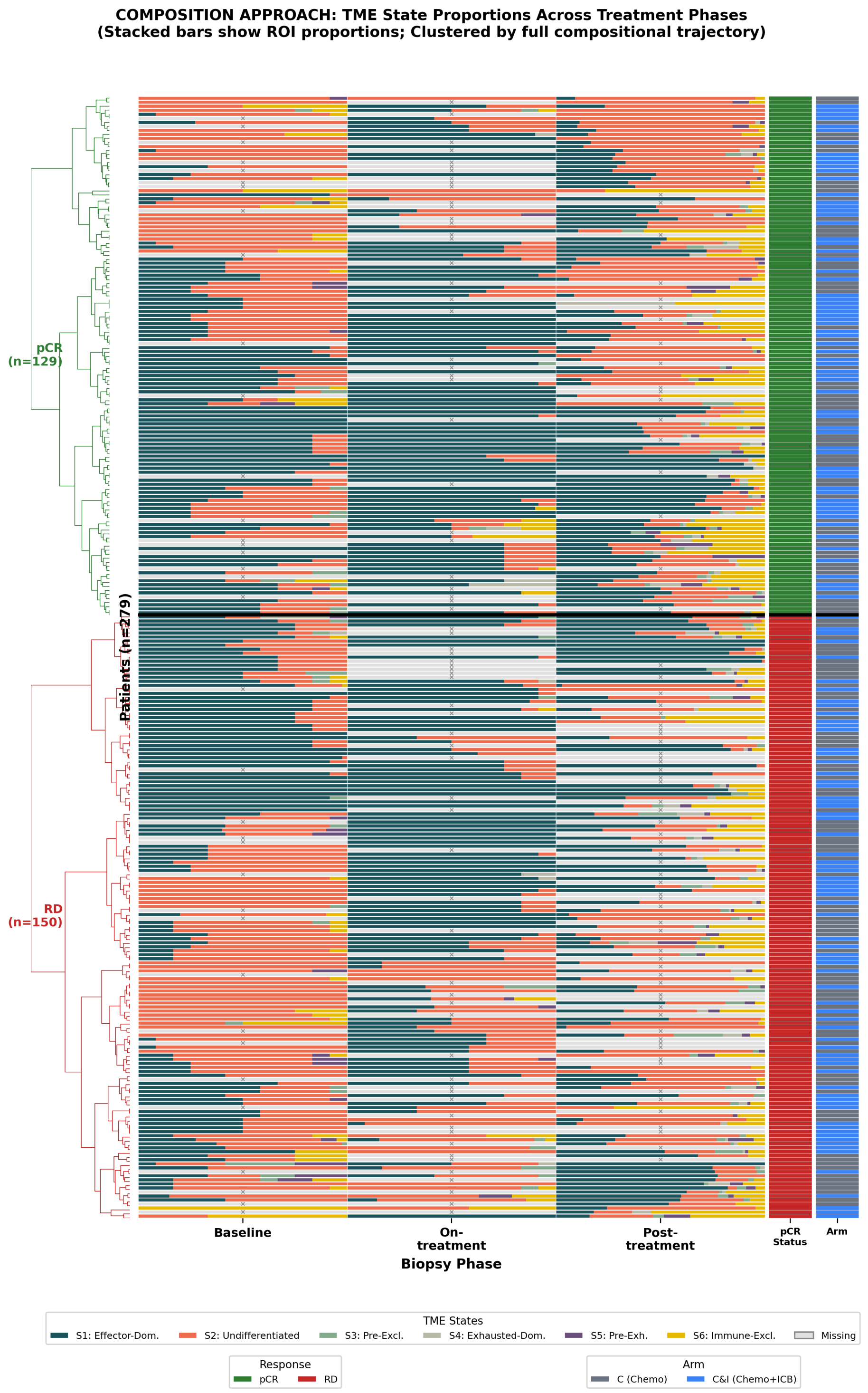


**Supplementary Figure 8. Patient trajectories in the IMC/Wang et al. data set.**

### Supplementary Figure 9

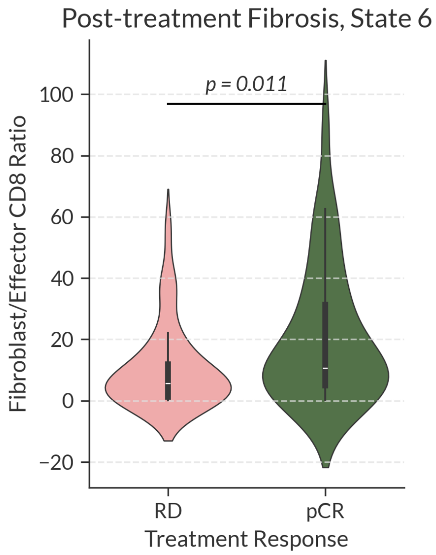


**Supplementary Figure 9. Fibroblast-to-Effector CD8+ T Cell ratio in TME State 6 patients at post-treatment**

#### Supplementary Figure 10

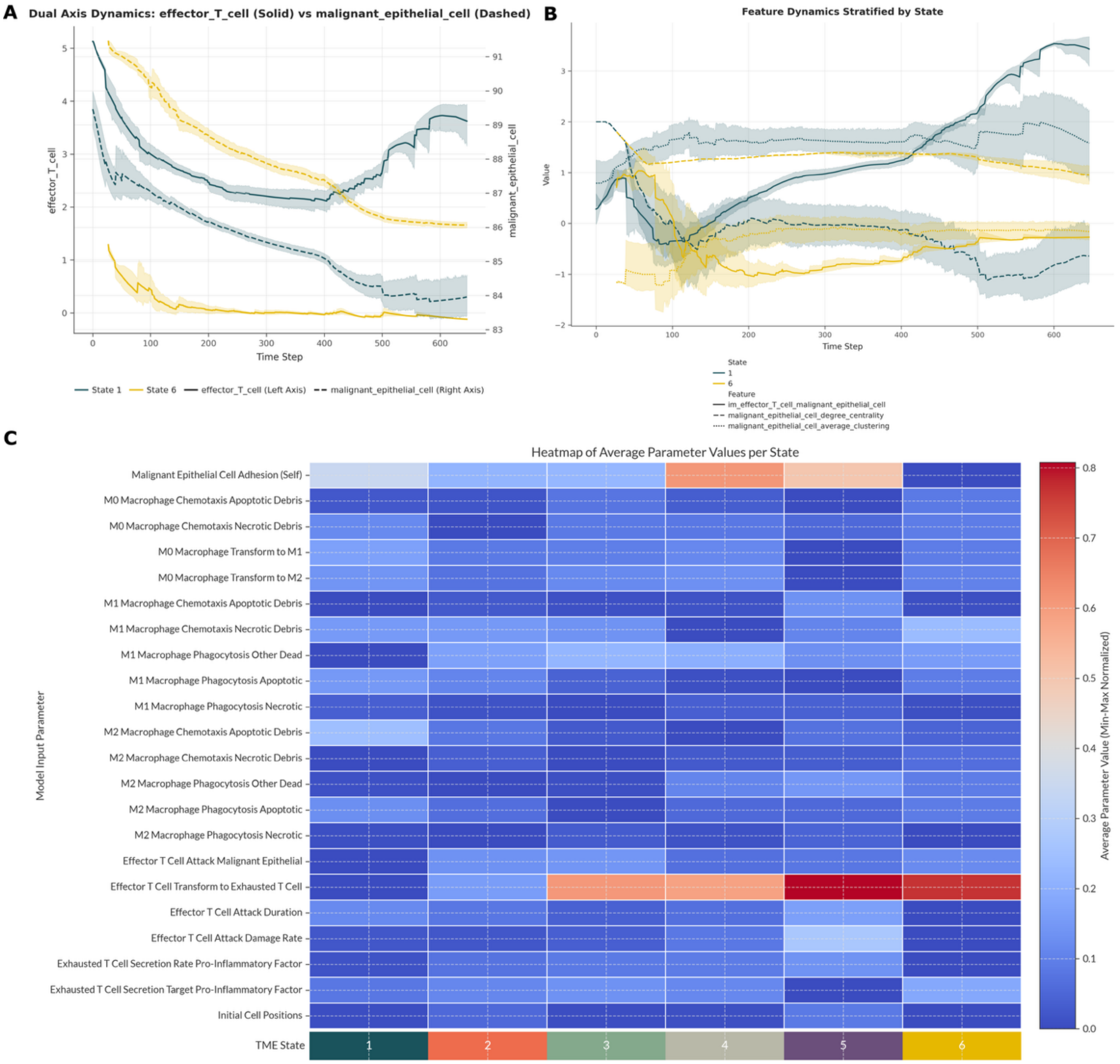


**Supplementary Figure 10. Analysis of within state dynamics and model input parameters. a,** Within-state population level dynamics. Left axis/solid line: effector CD8+ T cell population. Right axis/dashed line: malignant epithelial cell population. **b,** Within-state spatial dynamics (z-score normalized). Solid: effector–tumor interaction strength; dashed: tumor degree centrality; dotted: tumor clustering coefficient. **c,** Parameter associations with TME states. Heatmap shows mean parameter values (min-max normalized) across states.

#### Supplementary Figure 11

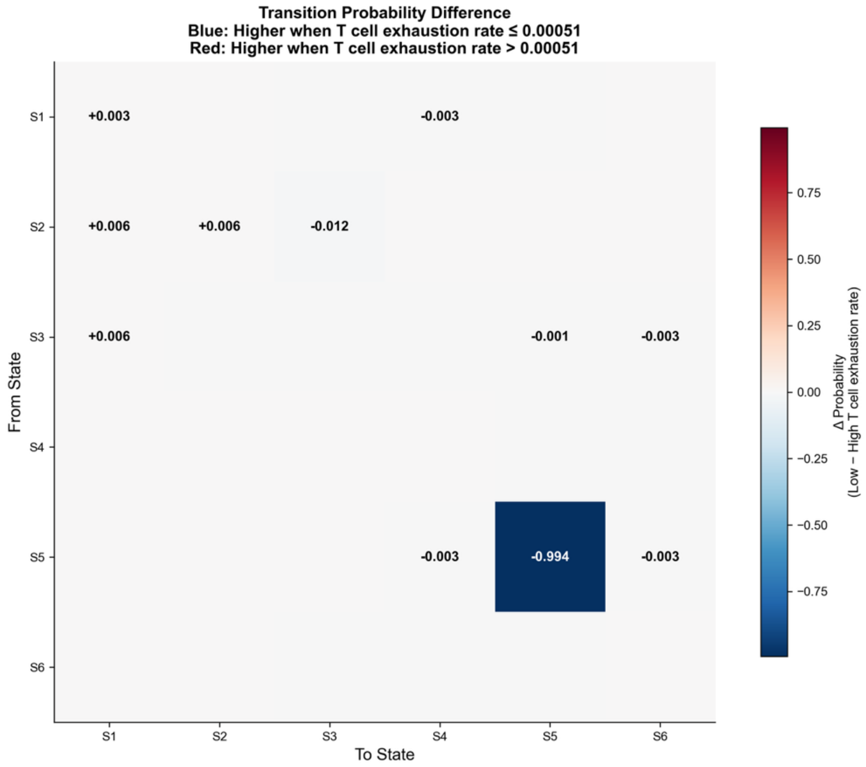


**Supplementary Figure 11. Transition Matrix Dynamics.** The difference in state transition probabilities conditioned on the value of the T cell exhaustion rate being set to the average value of simulation that terminated in the Effector-Dominated state.

#### Supplementary Figure 12

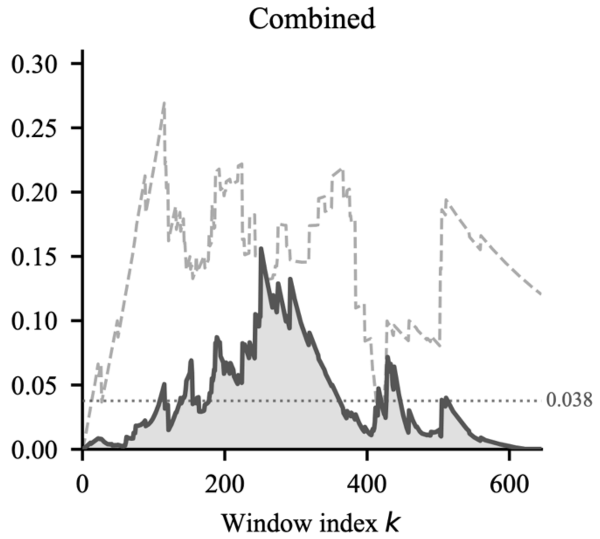


**Supplementary Figure 12. Kullback-Leibler (KL) Divergence Plot for Time-Varying Markov State Model (MSM).** The MSM is developed as a surrogate model for the ABM after defining the TME state landscape. The KL divergence measures the difference between the distribution of simulations occupying each state at each time point across the ABM simulations. The KL-divergence remains bounded by 0.16 with an average value of 0.038, indicating strong concordance between the ABM and MSM, and empirically supporting the use of the MSM as a surrogate to the ABM.

#### Supplementary Table 1: ABM Input Parameters

**Table S1.** Agent-based model parameters governing immune cell behavior and tumor-immune interactions. Parameter ranges were sampled using Latin Hypercube Sampling. All rates follow PhysiCell convention (1/min); adhesion strength and chemotactic sensitivity are dimensionless as debris signals are normalized.

| Parameter Name | Minimum Value | Maximum Value | Units | Citation |
|---|---|---|---|---|
| Malignant Epithelial Cell Adhesion Strength | 0.01 | 5 | dimensionless | [1,2] |
| M0 Macrophage Differentiation Rate to M1 & M2 | $2.3\times10^{-4}$ | $6.9\times10^{-4}$ | $min^{-1}$ | [3] |
| M0, M1 & M2 Macrophage Chemotactic Sensitivity to Apoptotic & Necrotic Debris | 0.01 | 1.0 | dimensionless | |
| M1 & M2 Phagocytosis Rate (apoptotic, necrotic, & other) | 0.001 | 1 | $min^{-1}$ | [4,5] |
| Effector T Cell Attack Rate | 0.05 | 5 | $min^{-1}$ | [6] |
| Effector T Cell Attack Duration | 0.1 | 10 | min | [7,8] |
| Effector T Cell Attack Damage Rate | 0.1 | 5 | $min^{-1}$ | [6] |
| Effector T Cell Exhaustion Rate | $4.9\times10^{-5}$ | $2.7\times10^{-3}$ | $min^{-1}$ | [9,10] |
| Exhausted T Cell Pro-inflammatory Factor Secretion Rate | 0.001 | 1.0 | $min^{-1}$ | [9,10] |
| Exhausted T Cell Pro-inflammatory Factor Environmental Target | 0.001 | 1.0 | dimensionless | [9–11] |

#### Notes on Units

**Dimensionless parameters:**

- *Adhesion strength*: Expressed as multiple of drag coefficient ν; values >1 indicate stronger adhesion than baseline drag
- *Chemotactic sensitivity*: Normalized because debris signal is treated as dimensionless (0–1 scale)
- *Environmental target*: Saturation density normalized to match pro-inflammatory factor substrate definition

**Rate parameters ($min^{-1}$):**

- *Differentiation rate*: Corresponds to ~24–72 hour differentiation timescale (1/rate = mean time to transition)
- *Exhaustion rate*: Corresponds to ~6–340 hour exhaustion timescale
- *Phagocytosis rate*: Rate per cell per minute of engulfment events
- *Attack/damage rates*: Per-minute probability or accumulation rate during cytotoxic contact

**Time parameters (min):**

*Attack duration*: Window of sustained T cell-target interaction
